# Transposable elements underlie chromosomal fusions and fissions in a highly species-rich group of butterflies

**DOI:** 10.64898/2026.06.15.732387

**Authors:** Camille Cornet, Tobias Baril, Charlotte Wright, Kay Lucek

## Abstract

Chromosomal fusions and fissions reshape karyotypes, recombination landscapes and patterns of speciation, yet the molecular mechanisms underlying their formation remain poorly understood. Comparing 37 chromosome-level *Erebia* genomes, a butterfly genus with exceptionally high rates of chromosomal rearrangements, we identify more than 250 fusion and fission events and characterise over one hundred breakpoints. Breakpoints and homologous regions in the most closely related species with the unfused chromosomal state are significantly enriched for repetitive elements, particularly R1-like LINE retrotransposons. This provides evidence for the implication of a specific LINE family in inter-chromosomal rearrangements that promote species diversification. R1-like elements at breakpoints are longer than copies in other genomic regions, consistent with ectopic recombination requiring sufficient sequence length and similarity. However, the burst of rearrangements in the youngest and most species-rich *Erebia* clade does not coincide with increased R1-like activity, indicating that repeat dynamics does not solely account for the elevated rates of fusion and fission. Indeed, we detect lineage-specific gains and losses of genes involved in DNA repair and chromatin organisation that coincide with this burst, suggesting a genomic context that facilitates chromosomal fusions and fissions. Our findings refine the role of repetitive elements in inter-chromosomal rearrangements, identify a candidate substrate for ectopic recombination in Lepidoptera, and establish a framework for understanding how karyotypic diversity arises.

## Introduction

Large-scale chromosomal rearrangements, such as fusions and fissions of chromosomes, are predicted to have significant evolutionary consequences both at micro- (*1*) and macroevolutionary scales (*2*). By altering the number, size and structure of chromosomes, these rearrangements can for example modify recombination landscapes and patterns of genetic diversity (*3*, *4*). Reduced recombination rates on fused chromosomes may also promote speciation by limiting gene flow between lineages with different karyotypes (*5*). Chromosomal fusions can also facilitate adaptation by linking adaptive alleles (*6*) or modulating gene expression (*7*). At a macroevolutionary scale, elevated rates of karyotypic change are often associated with increased rates of speciation (*8–10*). Strikingly, karyotype diversity varies widely across the Tree of Life, with some lineages displaying high karyotypic variation while others have conserved karyotypes over long evolutionary timescales (*11*, *12*). For example, mammals exhibit much higher rates of interchromosomal rearrangements than birds and reptiles (*13*), and clitellates display exceptionally scrambled genomes compared to other annelids (*14–16*). Given the evolutionary significance of chromosomal fusions and fissions, it is crucial to investigate the drivers of sudden shifts in chromosome evolution. In particular, the genomic features that underlie fusion and fission events remain poorly understood and may themselves represent important drivers of these shifts in rearrangement rates.

A potential source of variation in chromosome evolution is centromere organisation. The most extensively studied chromosomal fusions and fissions are known as Robertsonian rearrangements, which occur at, or close to the centromere (*17*, *18*). However, about 10% of all eukaryotic species have holocentric chromosomes that lack a single centromeric region but instead show centromeric activity scattered along the length of each chromosome (*19*, *20*). Novel chromosomal fusions and fissions are predicted to be more likely retained in holocentric species because meiotic defects in chromosomal hybrids may be less deleterious (*21–23*). The most species-rich holocentric clade is Lepidoptera, encompassing more than 160,000 butterfly and moth species (*24*, *25*). Despite holocentricity, most lineages have conserved the ancestral set of lepidopteran linkage groups, termed Merian elements (*26*). However, some clades have undergone massive bursts of chromosomal rearrangements, often associated with increased rates of speciation (*27*). One such clade is the genus *Erebia*, encompassing around 100 described species (*28*) that are often alpine and cold-adapted (*29*). *Erebia* species often differ markedly in their karyotypes, even between closely related species, with chromosome numbers ranging from 7 to 51 chromosome pairs (*8*, *30*). Even within *Erebia*, chromosome number changes are unequally distributed, with early-diverging clades having conserved chromosome numbers close to the modal *n* = 29. In contrast, the youngest and most species-rich *tyndarus* clade has evolved the highest karyotypic diversity (*n* = 8 – 51) over only the last 2.4 Myr (*8*). Intraspecific karyotype diversity has only recently been described for *Erebia* (*31*), but its extent throughout the genus is unknown.

A major gap in our understanding of the evolution of chromosomal fusions and fissions is their underlying genomic mechanisms (*32*). Repetitive genomic regions, such as transposable elements (TEs) or tandem repeats, have been suggested to cause rearrangements through ectopic recombination, whereby non-allelic repeats are recognised as homologous and recombine (*33*). This mechanism has been well described for intrachromosomal rearrangements, such as inversions in *Drosophila* (*34*, *35*) or small structural variants in humans (*36*). Inversion breakpoints in other species are often enriched for repetitive elements, consistent with ectopic recombination (*37*, *38*). However, evidence that ectopic recombination between repeats drives chromosomal fusions and fissions remains scarce. Tandem repeats have only recently been confirmed to underlie Robertsonian fusions in humans (*39*). Centromeric arrays of tandem repeats are also responsible for chromosomal fusions in holocentric plants (*40*, *41*). However, this centric model of chromosomal rearrangements cannot apply to Lepidoptera because their holocentromeres are not repeat-based (*19*, *42*). This suggests that other types of repeats may drive ectopic recombination associated with chromosomal fusions and fissions in Lepidoptera.

A major reason why the genomic drivers of chromosomal fusions and fissions remain largely unexplored is the lack of genomic resources needed to characterize breakpoints resulting from interchromosomal rearrangements in understudied species (*2*, *43*). The advent of long-read sequencing, enabling chromosome-level assemblies and improving the resolution of repetitive regions, now allows us to test the hypothesis that repeats underlie these rearrangements. Here, we use comparative genomics to identify the genomic features underlying chromosomal fusions and fissions across the butterfly genus *Erebia*, integrating a phylogenetic framework based on 37 chromosome-level reference genomes with comparisons between pairs of closely related taxa. First, we infer all fusion and fission events along the time-calibrated *Erebia* phylogeny from (*8*). We then test whether chromosomal rearrangements are phylogenetically associated with genome-wide proportions of repetitive elements and, if so, which repeat families are involved. Finally, we characterise chromosomal fusion and fission breakpoints, as well as homologous regions, between closely related species and taxa that do not carry the rearrangement, to test for repeat enrichment and determine whether specific repeat families underlie chromosomal fusions and fissions. We thereby address two related but distinct questions: (i) whether repetitive elements are associated with chromosomal fusions and fissions in *Erebia* butterflies and if so, (ii) whether repeat expansions can explain bursts of chromosomal rearrangements along the phylogeny.

## Results

### Genome assembly and annotation of *Erebia* butterflies

To characterize chromosomal rearrangements in *Erebia*, we combined 25 public chromosome-level reference genome assemblies with 12 additional genome assemblies from 11 species that we generated (Tables S1–S3). The complete dataset encompassed 36 species of the roughly hundred described in this genus, with a focus on the karyotypically highly diverse *tyndarus* clade (*8*). Assembly size ranged from 448.9 Mb to 647.5 Mb (Tables S2 & S3) and was not correlated with chromosome number (phylogenetic generalized least square regression (PGLS): *t*_1,35_ = 0.50, *p* = 0.620; Fig. S1A). The quality of all assemblies reached Earth Biogenome Project (EBP) standards (*44*), with contig N50 ranging from 4.2 Mb to 45.1 Mb, single-copy BUSCO scores ranging from 93.7 to 98.8%, k-mer completeness ranging from 87.5 to 99.7% and false duplication rate ranging from 0.18 to 3.10% (Tables S2 & S3). For the 12 assemblies that we generated, long-read RNA-seq data was available, showing mapping rates ranging from 93.5% to 99.9% (Table S2). The number of predicted genes in these assemblies ranged from 12’367 to 13’511, for 15’311 to 16’766 transcripts, respectively. Annotations reached single-copy BUSCO scores ranging from 92.1% to 95.1% (Table S2). For the other assemblies, for which RNA-seq data was not available, the number of predicted genes ranged from 16’960 to 36’935, for 19’037 to 41’049 transcripts, with single-copy annotation BUSCO scores ranging from 81.3% to 96.0% (Table S3).

We built a single *Erebia* repeat library iteratively, adding consensus sequences species by species, starting with the earliest-diverging ones in the time-calibrated phylogeny from (*8*) (see Methods). We then annotated each genome assembly with this single library and found that genome-wide repeat proportion varied from 53.5% to 63.8% and was correlated with assembly size (PGLS: *t*_1,35_ = 7.41, *p* < 0.001), but not with chromosome number (PGLS: *t*_1,35_ = 0.48, *p* = 0.638) (Fig. S1B-C). The repeat landscapes of all species were dominated by long interspersed nuclear elements (LINEs, 18.7% – 27.6% genomic proportion, the most common being L2), followed by rolling circles (also termed *Helitrons*, 12.9% – 17.9%; Fig. S2; Table S4).

### Chromosomal rearrangements along the *Erebia* phylogeny

We parsimoniously investigated the extent and phylogenetic distribution of chromosomal rearrangements across our available *Erebia* genomes and identified 182 fusion and 81 fission events across the genus (Figs. 1 & 2, Table S5). Two additional fusions of ancestral lepidopteran linkage groups, i.e. Merian elements (*26*), occurred prior to the origin of *Erebia*, leading from the modal chromosome number of Lepidoptera (*n* = 31) to that of *Erebia* (*n* = 29) (Fig. 2). Of the rearrangements occurring within *Erebia*, 24.7% of all fusions (n = 45) and 80.2% of all fissions (n = 65) occurred at the branch leading to, or within, the *tyndarus* clade, reaching rates of 12.2 fusions and 18.8 fissions per Myr, respectively (Fig. 1; Table S6). We found that two sequenced individuals were heterozygous for fusions, with two fusions in *E. gorge* (chromosomes 1 and 3) and one in *E. stirius* (chromosome 3) (Fig. S3), consistent with previous reports (*31*). Genome-wide repeat proportion was not associated with the number of fusions (PGLS: *t*_1,35_ = −1.69, *p* = 0.099) or fissions (PGLS: *t*_1,35_ = −0.29, *p* = 0.770) along the branches leading to extant species (Fig. S1D-E).

**Fig. 1.**
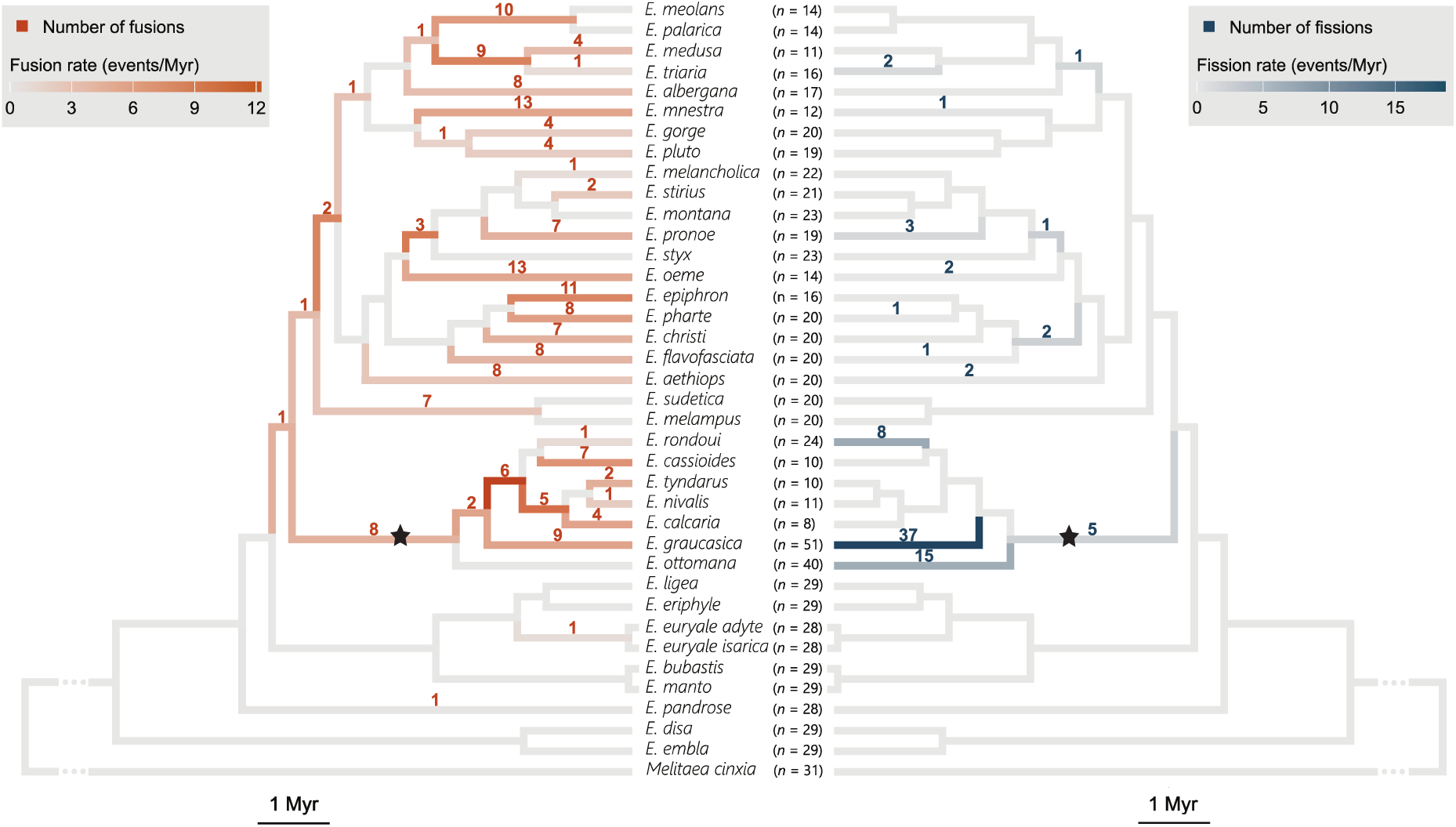
Chromosomal fusions (left) and fissions (right) along the time-calibrated *Erebia* phylogeny. Number of rearrangements are presented above each branch, and rates of rearrangements in terms of number of events per Myr are represented as colour gradients for each branch. The star represents the branch of the *tyndarus* clade, i.e. the youngest, most species rich and karyotypically diverse *Erebia* clade. Haploid karyotype is indicated for each species, excluding the W chromosome.

**Fig. 2.**
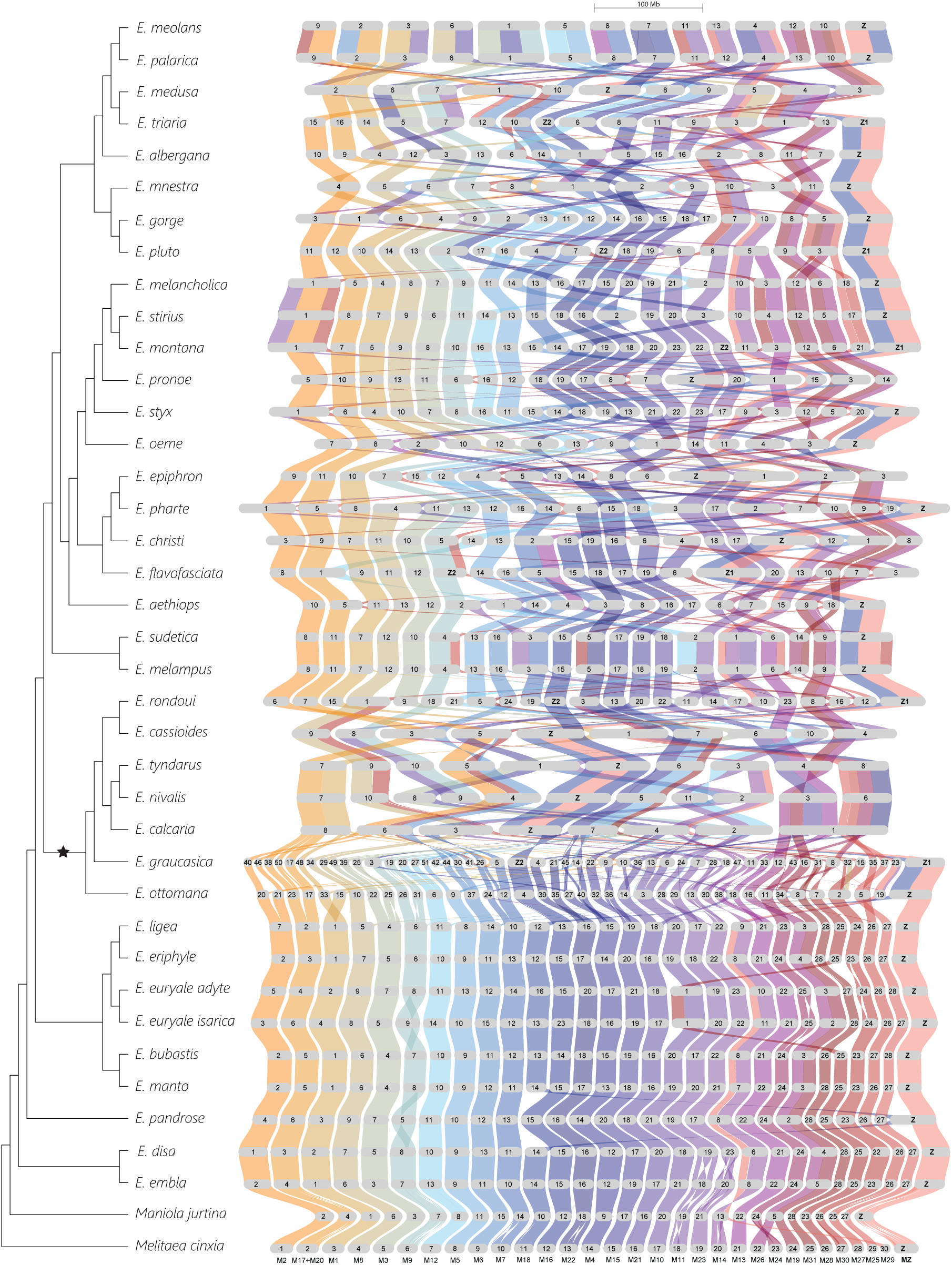
Synteny analysis showing homologous chromosomes and chromosomal rearrangements within the *Erebia* genus. Each grey rounded rectangle is a chromosome in an extant species and colours link ancestral chromosomes to their homologues in extant species. Chromosomes are scaled by size. Merian elements, ancestral linkage groups in Lepidoptera, are denoted with M in the outgroup *Melitaea cinxia*. The Z chromosomes are indicated in bold, and the W chromosomes are not shown. The phylogeny is presented as a cladogram and the star represents the *tyndarus* clade, the youngest, most species rich and karyotypically diverse clade in *Erebia*.

The Z chromosome did not undergo fissions, even in the species with the highest number of chromosomes, i.e. *E. graucasica* and *E. ottomana*, in which all ancestral autosomes underwent at least one fission event (Fig. 2). Ten independent cases of Z-autosome fusion involving the ancestral Z chromosome (Z1) were observed, each of which resulted in a neo-sex chromosome (Fig. 2 & S4A; Table S5). Six species had a Z1Z2W sex chromosome system (Tables S2 & S3), consistent with W-autosome fusions (*45*). Furthermore, the Z2 of *E. graucasica* (and putatively also *E. ottomana*) fused to the ancestral Z in the other species of the *tyndarus* clade, reverting the sex chromosome system to ZZ/ZW (Fig. 2). Some Merian elements were involved significantly more often than others in fusions (chi-square test: χ^2^ = 67.0, *p* < 0.001; Fig. S4A) and in fissions (χ^2^ = 100.3, *p* < 0.001; Fig. S4B), or were more likely to fuse with one another than others (Fig. S4C; Table S7). In particular, the neo-Z chromosomes repeatedly involved the same autosomes more often than expected by chance (10,000 permutations keeping the total number of fusions per Merian constant: Benjamini-Hochberg false discovery rate (FDR) adjusted *p* < 0.033; Fig. S4C; Table S7). The number of fusions in which a specific Merian element was involved was correlated with the number of fissions (Spearman correlation test: rho = 0.72, df = 29, *p* < 0.001; Fig. S4D). Taken together, these results suggest that the genomic regions undergoing rearrangements are not random.

To investigate whether bursts of chromosomal rearrangements are associated with repeat expansions, we tested the phylogenetic associations between the genomic proportion of each repeat family and chromosome number as well as the number of fusions or fissions along the branches leading to extant species. We only considered repeat families for which genomic proportion was > 0.01% in at least 20% of all species. Simple Repeats were significantly positively associated with the number of chromosomes (PGLS: *t*_1,35_ = 54.14, FDR adjusted *p* = 0.026; Fig. S1F; Table S8). Some DNA elements (TcMar-Fot1 and PiggyBac) as well as SINE/tRNA were significantly positively associated with the number of fissions (PGLS: all *t*_1,35_ > 11.5, all FDR adjusted *p* < 0.012; Fig. S1G-I; Table S8). Those three repeat families also showed a significant phylogenetic signal (all λ > 0.95, all FDR adjusted *p* < 0.003; Table S8). Although ancestral fissions were counted several times along the branches leading to the extant species, this should not affect our findings as most fissions occurred at the tips of the phylogeny (Fig. 1). We confirmed this by repeating the tests using only the number of fissions occurring at the tips, finding the same association with the three repeat families (Table S8). No repeat family was significantly associated with the number of fusions along the branches leading to extant species (Table S8). Given the high rate of chromosomal fusions and fissions in the *tyndarus* clade, we tested whether any repeat families had higher genomic proportions in this clade compared to other species. This was only the case for the DNA element TcMar-Fot1 (phylogenetic ANOVA with 10,000 simulations: *F* = 120.5, FDR adjusted *p* = 0.032; Fig. S1J; Table S8). We also tested if the *tyndarus* clade differed in repeat proportions compared to other *Erebia* clades (*8*). The only repeat that differed significantly between clades was DNA/CMC-EnSpm, for which genomic proportions were higher in the *embla* clade (phylogenetic ANOVA with 10,000 simulations: *F* = 87.1, FDR adjusted *p* = 0.005; Table S8). For the three repeat families that were associated with chromosomal fissions and with the *tyndarus* clade, we confirmed the presence of phylogenetic shifts in their genomic proportions at the base of the *tyndarus* clade using Bayesian reversible-jump multi-optima OU models (implemented in *bayou*; Fig. S5A-C; Table S9). Interestingly, Simple Repeats which were associated with chromosome number, and therefore indirectly the extent of fission, did not show any major shifts in the karyotypically variable *tyndarus* clade (Fig. S5D; Table S9).

Previous studies found that shorter Merian elements tend to be involved more often in chromosomal fusions and have higher repeat content than longer elements (*26*, *46*). We therefore asked whether the number of rearrangements in which a Merian element is involved in *Erebia* is associated with its sequence length or the proportion of specific repeat families. We used phylogenetic generalized linear mixed models, with Merian element as random factor. The number of fusions and the number of fissions were both significantly positively associated with the proportion of SINE/5S in each Merian (posterior mean slope = 2.70, 95% CI: 0.81 – 4.58, *pMCMC* = 0.006 and posterior mean slope = 7.80, 95% CI: 5.30 – 10.35, *pMCMC* < 0.001, respectively; Fig. S6A-B; Table S10). SINE/5S also showed a significant phylogenetic signal (λ = 1.00, FDR adjusted *p* < 0.001; Table S8). The associations of number of fusions and number of fissions with Merian size were not significant (posterior mean slope = 0.01, 95% CI: −0.05 – 0.06, *pMCMC* = 0.686 and posterior mean slope = −0.01, 95% CI: −0.10 – 0.08, *pMCMC* = 0.917, respectively; Fig. S6C-D; Table S10).

### Repeats underlie chromosomal fusion and fission breakpoints

We next tested for repeat enrichment at rearrangement breakpoints, focusing on the 111 autosomal rearrangements (47 fusions and 64 fissions) that occurred at the tips of the phylogeny and for which we had a closely related species for comparison that did not have the respective rearrangement (Table 1). We included the three fusions present in the heterozygous state of the phased reference genomes of *E. gorge* and *E. stirius* (Fig. S3; Table 1), representing likely recent rearrangements. We defined breakpoints as the interval between flanking blocks of syntenic orthologous genes with an average precision of 156 kb (± 23 kb SE) for fusions and 96 kb (± 13 kb SE) for fissions. We estimated enrichment by comparing breakpoints to 10,000 random genomic windows of the same size on the same chromosome, either in terms of the number of overlaps with each repeat family, or the total length of overlaps with each repeat family. Breakpoints for fusions are the regions resulting from the fusion, whereas breakpoints for fissions are the corresponding homologous region in the most closely related species retaining the unfissioned state. For fusions, we also examined the corresponding homologous regions in the most closely related species with the unfused state and for fissions, the two chromosome ends resulting from the fission. These homologous regions often corresponded to subtelomeric regions, in the pre-fusions or post-fission state (Table S11). We considered it *strong evidence* for a repeat association with chromosomal rearrangement when the fusion or fission breakpoint and both homologous regions were enriched for the same repeat family. We considered it as one of two levels of *supporting evidence* if either the breakpoint and one homologous region, or both homologous regions, were enriched for the same repeat family (see Fig. 3 & Fig. 4). We found strong evidence for 10% (11/111) of all breakpoints (13% and 8% of all fusions or fissions, respectively), supporting evidence from the breakpoint and one homologous region for 46% (51/111) of all breakpoints (62% and 34% for fusions and fissions, respectively) and from both homologous regions for 50% (55/111) of all breakpoints (36% and 59% for fusions and fissions, respectively; Table S11). Overall, 21% (10/47) fusions and 27% (17/64) of fissions did not show any statistical evidence for an association with repeats. Applying a Benjamini-Hochberg FDR correction, strong evidence remained for two fusions and two fissions, and supporting evidence for 21% (11/47) fusions and 20% (13/64) fissions (Table S11). To test for the role of segmental duplications, which have also been implicated in chromosomal rearrangements (e.g. (*47*, *48*)) we tested for each breakpoint whether the same segmental duplications were present at the breakpoint and at the corresponding homologous regions but found no such match.

**Fig. 3.**
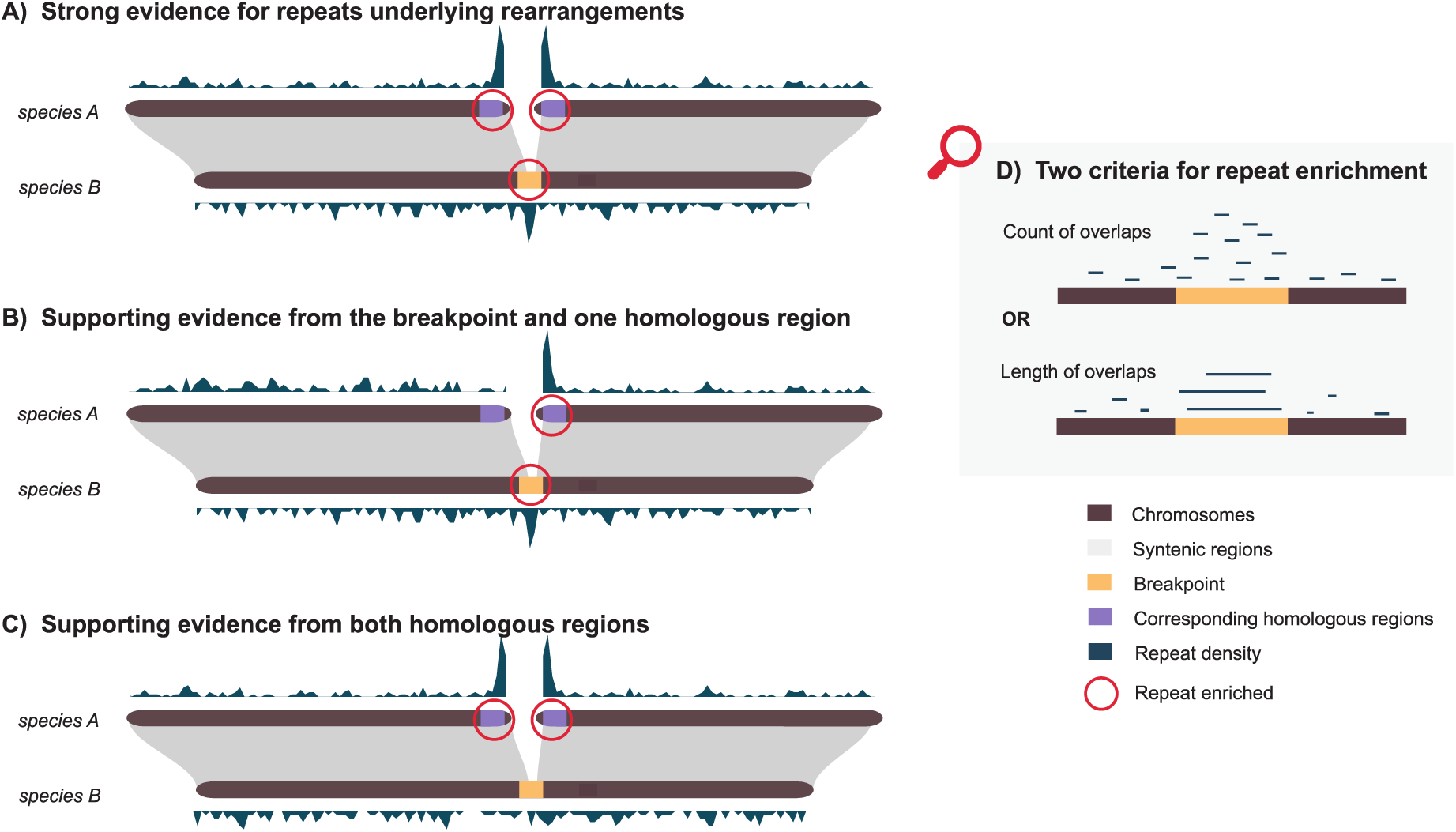
Schematic figure of the test for repeat enrichment at rearrangement breakpoints between closely related species A and B. For each fusion and fission breakpoint, we considered it strong evidence (A) for repeat association with the rearrangement if the breakpoint itself in species B and both corresponding homologous regions (in species A which has the unfused state) are enriched for the same repeat family. We consider it supporting evidence if either the breakpoint and one homologous (B) or both homologous regions (C) are enriched for the same repeat family. To be considered enriched in the same repeat family, the breakpoint and/or homologous regions need to be enriched for the same criterium, which is either the number of repeats overlapping with the breakpoint (or homologous region) or the total length of overlapping repeats (D). Note that repeats are not overlapping with each other in the dataset but are depicted as such in this figure for clarity.

**Fig. 4.**
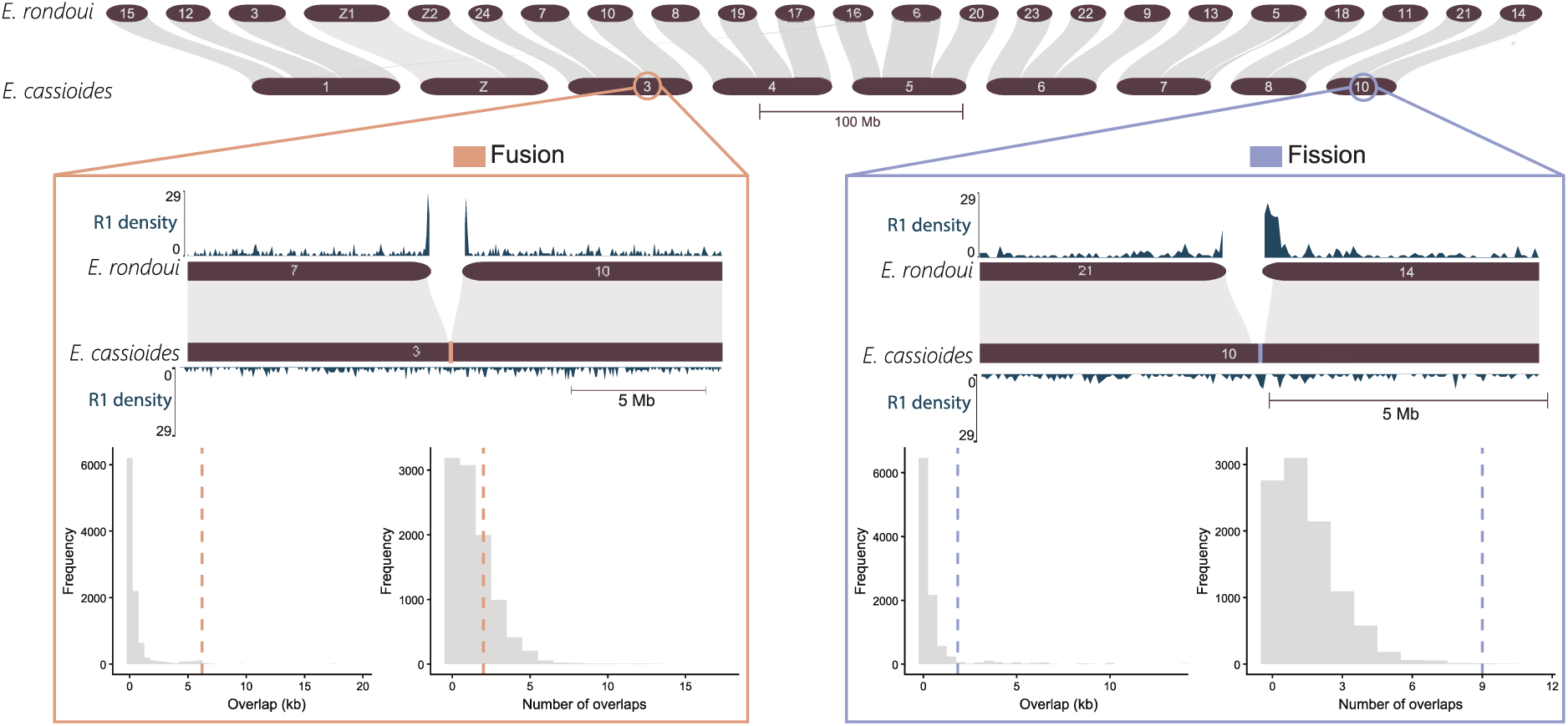
Example of a fusion and a fission breakpoint and associated enrichment of LINE R1-like elements for *E. rondoui* and *E. cassioides*. The density of R1-like elements was plotted using pyGenomeTracks v3.9 (*49*) with a window size of 50 kb. The histograms represent the results of the enrichment tests at the breakpoints in terms of lengths of overlaps (left) and number of overlaps (right). The grey distribution is the values at 10,000 random genomic windows; the dashed line is the value at the breakpoint. For the fusion, R1-like presents strong evidence for an involvement in the rearrangement before FDR correction and supporting evidence after correction. For the fission, LINE R1 presents strong evidence for an involvement in the rearrangement even after FDR correction. See Table S7 for details.

**Table 1.**
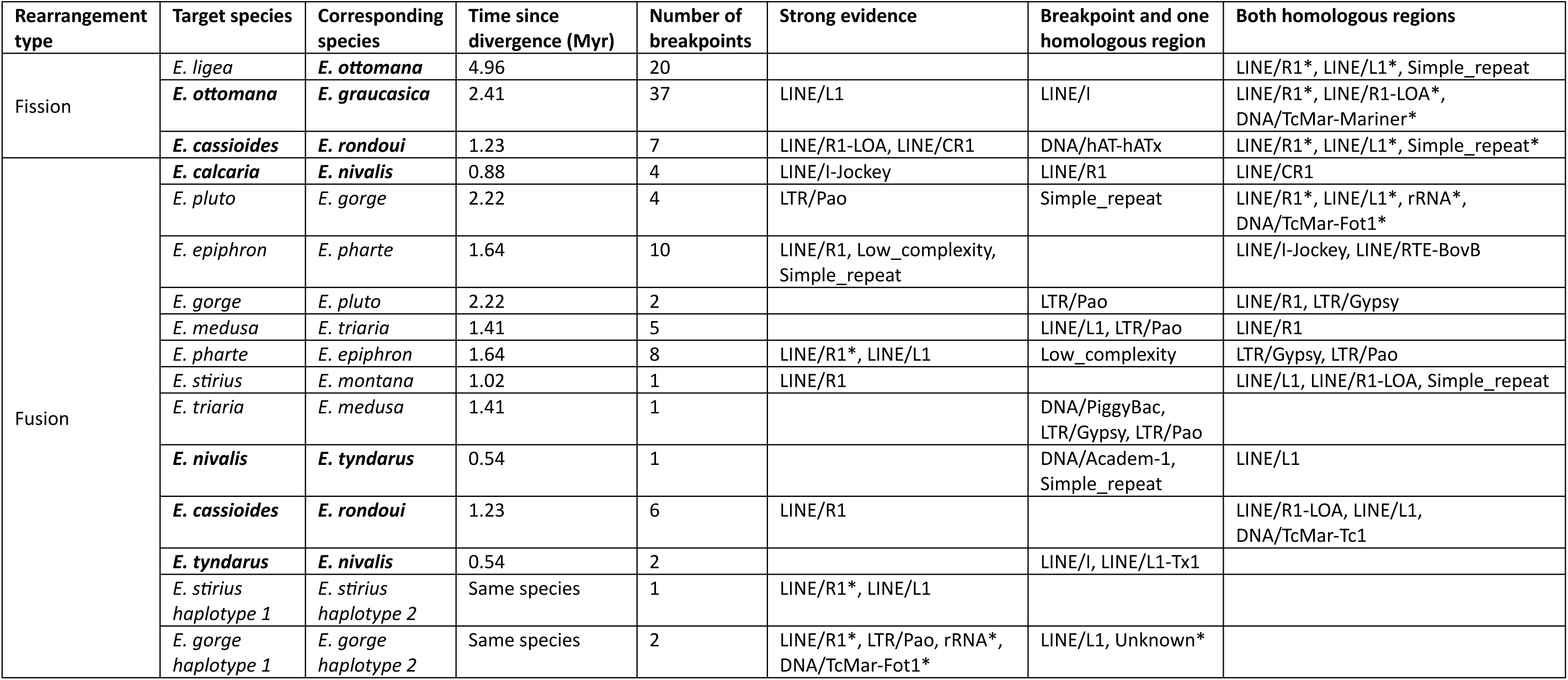
Repeat family enrichment tests at fusion and fission breakpoints and homologous regions in the most closely related species with the unfused state. Analyses with all breakpoints merged per species. For fusions, the target species is the one with the fusion; for fissions, the target species is the most closely related species with the fused state. Strong evidence means that a repeat family was enriched at the breakpoint and at both corresponding homologous regions. Supporting evidence means that this repeat was enriched in either breakpoint and one homologous region, or both homologous regions. Asterisks indicate repeats with the same level of evidence after Benjamini-Hochberg FDR correction. Species in bold belong to the *tyndarus* clade.

The repeats that were most often associated with both fusions and fission breakpoints were LINE R1-like elements, class I transposons of the Long Interspersed Nuclear Elements (LINEs) order that replicate using a “copy-paste” retrotransposition mechanism (*50*) (Table S11). R1-like elements represent between 0.55 and 1.26% of the genome of each species (Table S4). R1-like elements showed strong evidence for an association with 11% of all fusions (n = 5) and 2% of all fissions (n = 1), supporting evidence from the breakpoint and one homologous region for 15% of all fusions (n = 7) and for 2% of all fissions (n = 1), and supporting evidence from both homologous regions for 13% of all fusions (n = 6) and for 48% of all fissions (n = 31) (Fig. 5A-B, Table S11). A total of 29 other repeats were associated with rearrangements with strong or supporting evidence to a lesser extent, including LINE/R1-LOA, Simple Repeats and Low complexity regions (Fig. 5A-B, Table S11). The same repeats were frequently associated with both fusions and fissions. Only rRNA elements, which showed strong evidence in four fusions, were not associated with any fission. All repeats showing strong evidence for an association with fissions also showed at least supporting evidence for an association with fusions (Fig. 5A-B). Although the genomic proportions of the DNA elements TcMar-Fot1 and PiggyBac were significantly associated with the number of fissions along the phylogeny (Fig. S1G & S1H; Table S8), they only showed evidence for an association with two of our studied fission events (Fig. 5B, Table S11).

**Fig. 5.**
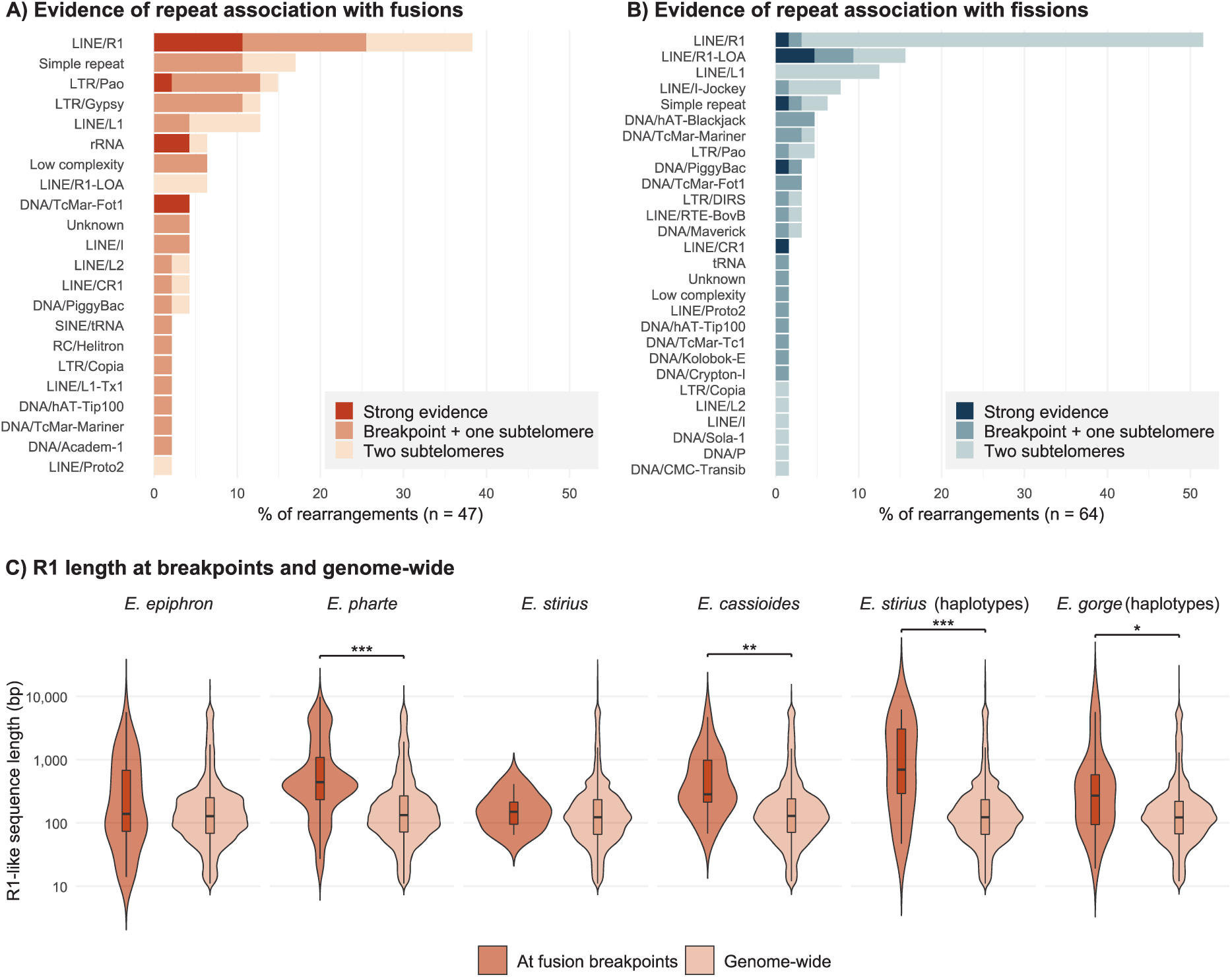
Number of fusions (A) and fissions (B) with which each repeat family is associated. Evidence of repeat association is divided into strong evidence, i.e. repeats enriched at the breakpoint and both corresponding homologous regions in a closely related species with the unfused state, or supporting evidence when repeats are enriched either at the breakpoint and one homologous region, or at both homologous regions. (C) Sequence length of R1-like elements at fusion breakpoints compared to genome-wide for the six species with strong evidence for R1-like elements association with fusions (permutation test: *** *p* < 0.001, ** *p* < 0.01, * *p* < 0.05).

To confirm the association of R1-like elements with rearrangements, we repeated the enrichment analysis using combined breakpoints for each species pair, rather than testing each breakpoint individually. Repeats associated with multiple rearrangements within the same species were expected to produce a stronger signal. We indeed found evidence for a LINE R1-like association with fusions for 8 out of 11 species pairs (4 with strong evidence, 4 with supporting evidence; Tables 1 & S12). For both species that were heterozygous for fusions, R1-like elements presented strong evidence for an association with fusions (Tables 1 & S12). For fissions, we found supporting evidence for an association with R1-like elements from both homologous regions in all three investigated species pairs (Tables 1 & S12). The association of R1-like elements with rearrangements presented above remained significant following an FDR correction (Tables 1, S11 & S12).

We further confirmed the importance of R1-like elements at chromosomal fusion and fission breakpoints using another approach that resamples random windows across the whole genome rather than per chromosome. We found consistent strong evidence for repeat association for 19% (9/47) fusions and 9% (6/64) fissions, supporting evidence involving the breakpoint and one homologous region for 53% (25/47) fusions and 34% (22/64) fissions, and involving both homologous regions for 43% (20/47) fusions and 49% (36/74) fissions (Table S13). The repeats most often associated with rearrangements were R1-like elements for 49% of all fusions and 55% of all fissions (15% and 3% with strong support, respectively; Fig. S7; Table S13). Combining breakpoints in each species pair, we found strong evidence for LINE R1-like association with fusions for 27% (3/11) of species pairs and in both species that were heterozygous for fusions, and supporting evidence for 27% (3/11) additional species pairs (Table S14). For fissions, there was supporting evidence for a LINE R1-like association in all 3 species pairs investigated.

Because ectopic recombination might be the underlying mechanism by which repeats facilitate rearrangement formation, and because recombination requires sufficient sequence length and sequence similarity (*51*), we compared the length and Kimura 2-parameter (K2P) distance distribution, a proxy for repeat divergence, between R1-like elements at fusion and fission breakpoints and genome-wide R1-like elements. For four of the six pairs showing strong evidence (36%, 4/11 of all pairs), and one species pair showing supporting evidence of association between R1-like elements and fusions (Table 1), the median length of R1-like elements was higher at breakpoints compared to their genome-wide median length (permutation test with 10,000 permutations: all *p* < 0.02; Figs. 5C & S8; Table S15). R1-like elements at breakpoints comprised 50 consensus sequences, of which we could orient 36 in the 5’ – 3’ direction, based on the presence of a canonical LINE poly-A tail at the 3’ end (or short tandem repeats other than poly-A), and 21 also possessed both canonical R1-like open-reading frames (*52*) (Fig. S9). Many R1-like elements at breakpoints were not truncated compared to the consensus, while some presented only the 3’ or 5’ end terminus (Fig. S9). R1-like elements did not show a different K2P distance at the breakpoints compared to the rest of the genome (permutation test with 10,000 permutations: all *p* > 0.065; Fig. S10; Table S15), except for two species for which R1-like elements at breakpoints had higher K2P distances (i.e. less sequence similarity: *E. calcaria* and *E. pharte*, *p* = 0.015 and *p* < 0.001, respectively; Fig. S10; Table S15). In addition, R1-like elements at breakpoints did not cluster together in our *Erebia* LINE phylogeny (Fig. S11). These results suggest that R1-like elements associated with rearrangements are longer but not more similar in sequence than those in the rest of the genome.

In many insects, LINEs play a role in telomere maintenance (*53*). R1-like elements may therefore be enriched at rearrangement breakpoints simply as a remnant of their telomeric and subtelomeric distribution, rather than reflecting an association with chromosomal rearrangements. To distinguish between these possibilities, we compared their genomic distribution to that of other repeat families. The distance of R1-like elements to chromosome ends was smaller than expected by chance (permutation test with 10,000 permutations: all *p* < 0.001; Fig. S12; Table S16), but this was also true for all repeats together (permutation test with 10,000 permutations: all *p* = 0.001; Table S16). Similarly, R1-like elements were not the only repeats enriched at subtelomeric regions compared to random genomic regions, in species used for the species pair comparisons (Fig. S13). Here, LINE/L1 elements were more enriched than R1-like elements at subtelomeric regions, both in terms of the number of overlaps (paired t-test: *t*_1,10_ = −6.12, *p* < 0.001) and the total length of overlaps (*t*_1,10_ = −2.50, *p* = 0.03), but did not show strong evidence for an association with chromosomal rearrangements (Fig. 5A-B). This suggests that R1-like enrichment at breakpoints and corresponding homologous regions cannot be solely explained by their distribution towards chromosome ends.

Despite LINE R1-like elements being often associated with chromosomal fusions and fissions in *Erebia*, the genomic proportion of R1-like elements showed no phylogenetic association with chromosome number, number of fusions or number of fissions along the phylogeny (PGLS: all *t*_1,35_ < 5.8, all *p* > 0.412; Table S8). The proportion of R1-like elements did not differ between the *tyndarus* clade and other species (phylogenetic ANOVA with 10,000 simulations: *F* = 0.09, *p* = 0.894; Fig. S10; Table S8), nor between the *tyndarus* clade and other clades (phylogenetic ANOVA with 10,000 simulations: *F* = 2.53, *p* = 0.851; Fig. S10; Table S8). A *bayou* analysis also did not show a shift in the proportion of R1-like elements at the branch leading to the *tyndarus* clade (Fig. S5E, Table S9). The Merian elements that underwent more fusions or fissions did not differ in terms of R1-like proportion (*MCMCglmm*: posterior mean slope = 0.08, 95% confidence interval: −0.44 – 0.59, *pMCMC* = 0.771 and posterior mean slope = 0.24, 95% confidence interval: −0.48 – 0.97, *pMCMC* = 0.505, respectively, Table S10). The average sequence length of R1-like elements did not differ in the *tyndarus* clade compared to other species (phylogenetic ANOVA: *F* = 3.25, *p* = 0.456) or clades (phylogenetic ANOVA: *F* = 2.95, *p* = 0.788). Mean sequence length of R1-like elements was also not associated with karyotype (PGLS: *t*_1,35_ = 0.60, *p* = 0.554), or the number of fusions (PGLS: *t*_1,35_ = 0.93, *p* = 0.357) or with fissions (PGLS: *t*_1,35_ = 0.80, *p* = 0.426) along the phylogeny. We further tested whether R1-like sequence similarity was higher in species within the *tyndarus* clade, finding that the mean K2P distance did not differ between R1-like elements in the *tyndarus* clade compared to other species (phylogenetic ANOVA: *F* = 0.98, *p* = 0.676; Fig. S10) or clades (phylogenetic ANOVA: *F* = 10.80, *p* = 0.133; Fig. S10). The mean K2P distance of R1-like elements was also not associated with karyotype (PGLS: *t*_1,35_ = 0.85, *p* = 0.400), or with the number of fusions (PGLS: *t*_1,35_ = −0.54, *p* = 0.590) or fissions (PGLS: *t*_1,35_ = 0.63, *p* = 0.536) along the phylogeny. In addition, R1-like elements in the *tyndarus* clade did not form a monophyletic cluster in the *Erebia* LINE phylogeny (Fig. S11) and the genomic distribution of R1-like elements did not differ between species of the *tyndarus* clade and other species (Fig. S12; Table S16).

### Gene family contractions and bursts of karyotypic evolution

We next asked whether gene family expansions and contractions could be associated with the increased rearrangement rate in the *tyndarus* group. We therefore performed a Gene Ontology (GO) enrichment test of gene family expansions and contractions along the *Erebia* phylogeny. We used non-RNA-seq-based gene predictions for all 37 *Erebia* genomes and identified 16’458 orthogroups at the root of the phylogeny, and an overall rate of gene gain and loss per gene family per million years (λ) of 0.035. Of these, 1’811 showed gene counts that changed significantly along the entire phylogeny. At the base of the *tyndarus* clade, 247 gene family expansions and 200 contractions occurred (Fig. S14A). We only considered GO terms enriched at the base of the *tyndarus* clade but not at the base of the karyotypically conserved *ligea* clade, to avoid including highly variable gene families. The 200 gene family contractions at the base of the *tyndarus* clade were significantly enriched for 56 Biological Processes GO terms (out of 27’186; Fig. S14B, Table S17). Among those GO terms were, notably, *non−recombinational interstrand cross−link repair* (6 genes lost) and *chromosome condensation* (20 genes lost). Gene family expansions in the *tyndarus* clade were significantly enriched for 68 GO terms (Fig. S14C, Table S18), of particular interest *rDNA chromatin condensation* (11 genes gained) and *UV-damage excision repair* (18 genes gained).

## Discussion

The mechanisms underlying chromosomal fusions and fissions, and the factors driving variation in chromosomal rearrangement rates across the Tree of Life, remain key open questions in chromosome evolution (*14*). Repetitive elements have been proposed as contributors to both (*54*, *55*), yet empirical evidence is often mixed (*56–58*). This is partly because many studies only focus on the repeat order level (*57*, *59*) when only some repeat families may cause rearrangements (e.g. (*35*)). Here, we built a fine-scale repeat library and explored the association between repeats and chromosomal rearrangements in the butterfly genus *Erebia*, first at a phylogenetic scale, correlating repeat dynamics with rearrangement rates. Second, at the level of individual recent breakpoints to investigate the genomic patterns underlying fusions and fissions. *Erebia* is one of the most karyotypically variable known butterfly genera as a result of chromosome fusions and fissions (*27*). This diversity in chromosome numbers and rearrangements is not equally distributed among the different subclades but is especially concentrated in the young and species-rich *tyndarus* clade (*8*, *30*). Using chromosome-level genome assemblies, we show that rates of fusions and fissions in the *tyndarus* clade are even higher than previously inferred using karyotypes (Fig. 1; (*8*)). For example, the sibling species *E. tyndarus* and *E. cassioides* that form narrow contact zones with few F1 hybrid males (*60*, *61*) share the same karyotype (*n* = 10), yet we identified seven distinct chromosomal fusions in each lineage since their divergence from a common ancestor (Fig. 1, Table S5). These lineage-specific rearrangements may contribute to the absence of gene flow between the two species (*61*). Across our studied *Erebia* species, we identified 182 fusion and 81 fission events that have occurred over the past 5.4 Myr (Figs 1 & 2). Most of these rearrangements, particularly fissions, occurred at the base of, or within, the *tyndarus* clade, strengthening previous evidence of accelerated karyotypic evolution in this clade (*27*, *30*), which is associated with elevated rates of speciation (*8*). Such a high rate of chromosomal fusions and fissions is rare among Lepidoptera, as most lineages have retained the ancestral karyotype (*26*, *27*). Using chromosome-level genome assemblies, similarly high rates of fusions and fissions have only been found in a few other butterfly clades, including Ithomiini (*62*) and *Leptidea* (*47*, *57*). Equally high rates of fusions and fissions occur in holocentric *Carex* sedges (*63*). Beyond holocentric species, even the most karyotypically variable mammal clades reach rates of chromosomal rearrangements that are ∼6 times lower than the rates observed in *Erebia* (*64*).

We observed that the Z chromosome in *Erebia* underwent ten independent fusion events to an autosome, forming neo-sex chromosomes (Figs. 2 & S4). This number of independent fusion events is lower than for fusions between autosomes. This pattern is consistent with observations in aphids (*56*) and other groups of butterflies with high rates of rearrangement (*65*), where the X or Z chromosome is less likely to undergo interchromosomal rearrangements compared to autosomes. However, it contrasts with more karyotypically conserved Lepidoptera clades, where the Z chromosome is the chromosome that is most frequently involved in chromosomal fusions (*26*). This might be due to a relatively old fusion between the Z chromosome and M18, shared by 78% of our *Erebia* species with subsequent Z-autosome fusions being rarer (Fig. 2). Notably, while all autosomes experienced fissions in *Erebia*, no fissions were detected for the Z chromosome, nor for the Merian element that became sex-linked through the early Z–autosome fusion (Figs. 2 & S4). A similar pattern has been reported for *Polyommatus atlanticus*, a butterfly with extensive chromosomal fissions, in which the two Z chromosomes are the only ones that did not undergo rearrangement (*58*). Together, these observations suggest that the relaxation of constraints on chromosomal evolution in *Erebia* does not apply uniformly across the genome, and that the Z chromosome remains subject to stronger constraints than autosomes. One possible explanation is stronger purifying selection acting on the Z chromosome due to its full exposure in heterogametic females (*66*), which may outweigh the effects of reduced effective population size on the Z chromosome. If most chromosomal fusions are slightly deleterious and reach fixation primarily through genetic drift, as proposed for other butterflies (*65*), fusions would be expected to fix more readily for autosomes than for the Z chromosome, where the strength of purifying selection likely outweighs that of drift. Additional constraints related to sex-chromosome-wide dosage compensation (*67*) or to pairing and segregation during meiosis (*68*) may limit the tolerance of fissions on the Z chromosome.

We found that many fusion and fission breakpoints are enriched for a single repeat family, R1-like elements (Fig. 5). By testing for repeat enrichment at breakpoints and their corresponding homologous regions in closely related species with the unfused state (Figs. 3 & 4), we mitigate a limitation of comparative genomics, i.e. that breakpoints are often highly repetitive and subject to rapid sequence turnover, leading to evolutionary noise (*32*). The R1-like enrichment is consistent with ectopic recombination, as observed for other LINE families. For example, LINE-1 is involved in translocations in humans (*69–71*). LINE enrichment was also found at fusion breakpoints in *Leptidea* butterflies (*57*) and in the silkworm *Bombyx mori* (*59*). However, as these analyses were done at the repeat order level, it remains unclear whether specific LINE families, such as R1-like elements, are specifically associated with rearrangements in these systems, also because LINEs are the most abundant transposable elements in Lepidoptera (*72*). At repeat order level, evidence for ectopic recombination through repeat enrichment at breakpoints is limited (*56–58*). Segmental duplications may also be associated with rearrangements, as has been found in mammals (*48*) and *Leptidea* butterflies (*47*). Focusing on breakpoints and homologous regions simultaneously, we did not find an effect of either tandem repeats or segmental duplications in *Erebia*.

Some R1-like elements, such as TRAS and SART (*73*) are known for their role in telomere maintenance in Lepidoptera (*74*). Therefore the R1-like elements that we found to be associated with chromosomal fissions might have the dual role of facilitating rearrangements through ectopic recombination while simultaneously stabilizing chromosome ends by elongating telomeres, which illustrates the balance between costs and benefits of TEs (*75*). Another possibility is that the presence of R1-like elements at breakpoints primarily represents remnants of repeats that were telomeric in species with the unfused state. This seems unlikely as most repeats are enriched at chromosome ends without showing an association with rearrangements in our analyses (Fig. S13; Table S16). A major question that arises from our results is why R1-like elements are more often associated with rearrangements than other LINEs. If canonical LINE tandem repeats at the 3’-end (e.g. poly-A (*76*)) served as the substrate for ectopic recombination, rearrangements should not be specific to R1-like elements, but instead involve other LINE families as well. Recombination requires a minimum sequence identity and length (e.g. 100 bp in yeast and 500-1000 bp in mammals (*51*, *77*)), and R1-like elements are longer at breakpoints where they are associated with rearrangements than in other regions of the genome (Fig. 5). We therefore hypothesize that other parts of the R1-like elements, potentially long and relatively conserved open reading frames (Fig. S9), might be the sequences recognized during recombination. This is consistent with the rate of ectopic recombination scaling with the length of non-LTR retrotransposons in *Drosophila* (*77*), and with the finding of longer LINE sequences at rearrangement breakpoints were also detected in mammals (*78*), and because longer homologous sequences increase the likelihood of recombination (*51*), this pattern is consistent with ectopic recombination as the underlying mechanism of rearrangements. A non-negligible proportion of rearrangements showed no statistical evidence for repeat involvement (Fig. 5). Although signals of ectopic recombination may have decayed over evolutionary time, alternative mechanisms, such as non-homologous end joining (*79*), replication-based processes (*80*), or LINE retrotransposition that directly bridges chromosomal breakpoints (*81*) may also account for the rearrangements lacking detectable repeat enrichment.

In Lepidoptera, the modal karyotype of *n* = 31 has been conserved for over 150 Myr, with chromosomal fusions and fissions being rare and largely restricted to a few clades (*24*, *26*, *27*). Within *Erebia*, we show that the *tyndarus* clade evolved dozens of chromosomal fusions and fissions in just ∼2.4 Myr (Fig. 1; (*8*)). Consistent with the hypothesis that this burst of rearrangements could be driven by expansions of repetitive elements, the genome-wide proportions of some families, rather than orders, of DNA elements were associated with the number of fissions along the *Erebia* phylogeny (Fig. S1 & S5). Merian elements undergoing more fusions and fissions were further enriched for 5S rRNA-derived SINEs (Fig. S6). A previous phylogenetic study suggested that rare *Helitron* elements were phylogenetically associated with chromosome fissions in *Erebia* (*54*). However, that study relied on low-coverage, reference-free short-read data, and we did not recover this signal using high-quality chromosome-level reference genomes and an iteratively built repeat library. Importantly, in the present study, the repeats enriched at fusion and fission breakpoints were not the same repeats showing a phylogenetic association with chromosomal rearrangements (Figs. 5 & S1). Although R1-like elements were frequently associated with chromosomal fusion and fission breakpoints, their genomic proportions and age distributions were not correlated with the number of rearrangements across the phylogeny, nor with the *tyndarus* clade specifically (Fig. S10). Evidence for repeats underlying fusions and fissions without a consistent phylogenetic pattern in changes of rates of rearrangements was similarly observed in turtles (*55*). While we cannot exclude the possibility that a burst of R1-like elements occurred during the diversification of the *tyndarus* clade that is no longer detectable, our results indicate that expansions of repetitive elements alone are unlikely to account for the burst of chromosomal rearrangements observed in this clade.

In *Erebia*, R1-like elements occur at similar genomic proportions in both karyotypically conserved, early-diverging species and species with highly rearranged genomes (Fig. S10), suggesting that TE activity alone does not drive the burst of chromosomal rearrangements. Instead, gains or losses of genes involved in maintaining genome stability, such as DNA repair, recombination, or telomere maintenance, could allow for the evolution of numerous chromosomal rearrangements (*82*). At the base of the *tyndarus* clade, gene family contractions and expansions were enriched for functions that could influence chromosomal rearrangements, either directly through DNA repair or indirectly via chromatin structure and mitotic/meiotic processes (Fig. S14, Tables S17 & S18). Although focusing on *Erebia* and given that GO term results should be interpreted cautiously (*83*), our findings highlight the possibility of a role of gene gains and losses in facilitating bursts of rearrangements. Indeed, losses of DNA repair genes have previously been associated with chromosomal fusions in birds (*84*) and fissions in mammals (*85*). Repeat activity could similarly account for such changes, as has been found in gibbons, where insertions of retrotransposons into chromosome segregation genes drives genome plasticity (*86*). We observed losses of genes involved in chromatin condensation, which may contribute to rearrangements given that chromosomal fissions in some Lepidoptera occur more frequently in open chromatin (*58*). In holocentric organisms, chromosomal fusions and fissions may be more readily tolerated than in monocentric organisms (*21–23*), potentially allowing changes in genes involved in genome stability to persist. Conversely, maintenance of genes involved in genome stability could explain why TE expansions are not accompanied by bursts of chromosomal rearrangements in some clades (*87*). Under this hypothesis, most species would retain conserved karyotypes while their DNA repair machinery is intact, whereas sudden bursts of chromosomal rearrangements could occur following the loss of key genome stability genes.

## Conclusions and future perspectives

The causes of variation in chromosomal rearrangement rates across taxa, and the mechanisms by which fusions and fissions arise, remain poorly understood. Here, we show that chromosomal fusions and fissions are even more prevalent in *Erebia* than previously inferred by karyotyping, especially in the *tyndarus* clade where most fissions occurred. These results strengthen previous findings, that found rates of chromosomal rearrangements to be associated with increased speciation rates in this clade (*8*). Fusion and fission breakpoints in *Erebia*, as well as the corresponding homologous regions in closely related species, are frequently enriched for R1-like elements. This pattern is consistent with evidence from other species where LINEs were enriched at the breakpoints of large-scale chromosomal rearrangements (e.g. (*69*)) and is compatible with ectopic recombination contributing to chromosomal evolution. However, expansions of R1-like elements are not associated with the overall burst of chromosomal rearrangements in the *tyndarus* clade, as R1-like abundance is similar across species, including those with the most conserved karyotypes. Instead, our data suggests that lineage-specific gains and/or losses of genes involved in genome stability, such as DNA repair or chromatin organization, may influence the probability that fusions and fissions arise, facilitating episodic bursts of chromosomal rearrangements. Chromatin context is also likely to influence rearrangement formation (*16*, *88*). With the growing availability of Lepidopteran genomes (*43*), our results establish a framework to test the genomic drivers of chromosomal rearrangements at macroevolutionary scale.

## Materials and Methods

### Butterfly sample collection

We collected adults for genome sequencing and annotation for 12 *Erebia* genomes from 11 species in the Alps, the Massif Central and the Caucasus during the flight seasons in 2021-2023 (Table S1). We included two subspecies of *E. euryale*, i.e. *adyte* and *isarica*, that show strong genomic differentiation (*89*). We chose those species and subspecies to cover a broad phylogenetic and karyotypic range, from *n* = 8 in *E. calcaria* to *n* = 51 in *E. graucasica* (*8*). Of the 12 samples, 11 were females, the heterogametic sex in Lepidoptera (*68*). We kept the specimens alive until storage at −80°C.

In addition to those 12 species, we included 27 publicly available chromosome-level reference genomes: two outgroups (*Melitaea cinxia* and *Maniola jurtina*) and three *Erebia* species (*E. ligea*, *E. aethiops*, *E. epiphron*) sequenced by the Darwin Tree of Life Project (*90*); *E. palarica* (*91*); and 21 *Erebia* species (*E. albergana*, *E. bubastis*, *E. christi*, *E. disa*, *E. embla*, *E. eriphyle*, *E. flavofasciata*, *E. gorge*, *E. manto*, *E. medusa*, *E. melampus*, *E. melancholica*, *E. meolans*, *E. mnestra*, *E. montana*, *E. pharte*, *E. pronoe*, *E. rondoui*, *E. stirius*, *E. sudetica* and *E. triaria*) produced by Project Psyche (*43*) (Table S3).

### Sequencing strategy

High-molecular weight DNA extraction and sequencing strategy followed (*92*): for genome assembly, we generated PacBio HiFi reads from half a thorax (average of 1.45 million HiFi reads or 12.9 Gb per sample; Table S1) to reach a target coverage > 25x. PacBio library preparation and sequencing on the Sequel IIe and Revio instruments were performed by the Genomics Technologies Facility (GTF, Lausanne, Switzerland). For assembly scaffolding, we generated Hi-C data using the heads of the same individuals that we used for PacBio sequencing with the Arima High Coverage HiC kit (Arima Genomics, San Diego, CA, USA). Illumina sequencing on the Novaseq 6000 instrument was performed by the Next Generation Sequencing Platform (NGS Platform, Bern, Switzerland), resulting in an average of 172 million 150 bp paired reads per sample (Table S1).

For the genome annotation of the 12 reference genomes generated here, we extracted RNA from half the thorax and the head of another individual from the same or nearby location (Table S1) with the Qiagen RNeasy Plus Universal Mini Kit (Qiagen, Hombrechtikon, Switzerland). Following PacBio Kinnex multiplexing and Iso-Seq sequencing on one PacBio Revio SMRT cell (NGS Platform, Bern, Switzerland), we obtained on average 11.40 million HiFi reads or 17.4 Gb per sample (Table S1), aiming to obtain full-length transcripts.

### Genome assembly

The genome assembly pipeline for all species that we collected followed (*92*). Briefly, for each sample, we assembled the HiFi reads into contigs using hifiasm v0.16.0-r369 (*93*) in primary assembly mode. We then used purgedups v1.2.5 (*94*) to remove haplotypic duplication from the primary assembly. For scaffolding of the draft assembly into chromosomes, we mapped the Hi-C reads with the Arima mapping pipeline v03 (https://github.com/ArimaGenomics/mapping_pipeline) and scaffolded the assembly using YaHS v1.2 (*95*), followed by manual curation using Juicebox v1.11.08 (*96*). We assembled the mitogenomes from the raw HiFi reads using MitoHiFi v3.0.0 (*97*). We manually removed scaffolds corresponding to haplotypic duplications or unplaced repetitive regions to reach chromosome-level primary genome assemblies (see final Hi-C maps in Fig. S15). To confirm that the final assemblies reach the Earth Biogenome Project (EBP) standards (*44*), we assessed their quality in terms of gene content and k-mer completeness, contiguity, base pair quality and false duplication rates. We identified the Z and W sex chromosomes (Fig. S16) based on the coverage of the HiFi reads (for females) and Illumina reads (for males, data from (*8*)) mapped back to the assembly, except for *E. ottomana* for which we sequenced a male and identified the Z chromosome by synteny with Merian element Z.

### Genome annotation

For our genome annotations, we first built a repetitive elements library for *Erebia*, selecting 14 (sub)species spanning a broad phylogenetic and karyotypic range (*E. pandrose*, *E. euryale adyte*, *E. euryale isarica*, *E. ligea*, *E. ottomana*, *E. graucasica*, *E. calcaria*, *E. nivalis*, *E. cassioides*, *E. aethiops*, *E. oeme*, *E. styx*, *E. pluto*). We started by detecting repetitive elements *de novo* in the earliest-diverging species, i.e. *E. pandrose*, using EarlGrey v3.1 (*98*) configured with the Dfam v3.7 curated set of repeats for Arthropoda (*99*). We then used the output library as a starting point for the next earliest-diverging species, moving along the phylogeny, to get a broad and representative repeat library for the genus. We clustered the resulting library to retain repeats with a maximum of 80% sequence identity in at least 80% of their aligned sequences using cd-hit-est v4.8.1 (*100*), satisfying the 80-80-80 rule (*76*, *101*). To ensure that our library did not contain any host genes (e.g. genes from duplicated butterfly gene families), we used diamond blastx v2.0.15.153 (*102*) with the Arthropoda NR database (downloaded on 18.09.2024) and the repeat library as query. Matching sequences were then submitted to CD-Search (https://www.ncbi.nlm.nih.gov/Structure/bwrpsb/bwrpsb.cgi, accessed on 24.09.2024). Repeat sequences that matched the Arthropoda NR database but returned no CD-Search hit to a known TE-associated protein domain (following (*76*)) were considered as host genes and removed from the final *Erebia* repeat library. This conservative approach ensures that no conserved host gene is masked prior to gene prediction.

We softmasked all *Erebia* genomes with the final *Erebia* repeat library using RepeatMasker v4.1.5 (*103*) and the rmblast search engine. For the two outgroups, *Maniola jurtina* and *Melitaea cinxia*, we used the Arthropoda Dfam v3.7 database instead of the *Erebia* repeat library. We then used a different gene prediction approach for the 12 genomes that we assembled and for the publicly available genomes. For the former we assessed the quality of the HiFi reads from RNA sequencing using NanoPlot v1.32.1 (*104*) and mapped the raw HiFi reads to the genomes using minimap2 v2.21 (*105*) with the splice:hq preset. We used the resulting alignment files as well as the Arthropoda OrthoDB v11 protein set (*106*) as evidence for BRAKER3 v3.0.8 (*107*), with the long read RNA-seq and protein data mode. For the other genomes, for which no RNA-seq was available, we used BRAKER2 v3.0.7 (*108*) with the evidence from the Arthropoda OrthoDB v11 protein set. For all genomes, after filtering the gene set to retain only the longest transcript per gene, we assessed genome annotation quality using BUSCO v5.7.1 (*109*) with the Lepidoptera OrthoDB v10 in transcriptome mode. We then functionally annotated the genomes using Fantasia v3.0.0 (*110*) with the protein language model ESM3c to get Gene Ontology (GO) term assignments for each gene.

### Macrosynteny and correlations between repetitive elements and chromosomal rearrangements

To detect large-scale chromosomal rearrangements along the phylogeny between the 37 *Erebia* reference genomes, we first ran OrthoFinder v2.5.5 (*111*) on the predicted gene sets restricted to the longest isoform per gene to identify orthologous and paralogous genes across the *Erebia* phylogeny (*8*). We then identified and plotted synteny blocks across the genus with GENESPACE v1.3.1 (*112*) based on the OrthoFinder run described above and using blkRadius = 50 and blkSize = 5. We also ran syngraph v0.1.0a (*113*), which identifies chromosomal fusion and fission events along each branch of the phylogeny using a parsimony approach based on the genomic position of the orthogroups defined by OrthoFinder. We excluded translocations as they are impossible to distinguish from a succession of fissions and fusions, and because syngraph may lead to false positive translocation detection (*113*). Syngraph is conservative as it does not account for intrachromosomal rearrangements. In three cases, syngraph inferred a fission event of the Z chromosome followed by a fusion event, as it was equally parsimonious as two independent fusions of the Z to autosomes. Because there is no known fission event involving the Z chromosome in Lepidoptera (*26*), we considered two independent fusions as more likely. We calculated rates of fusions and fissions in terms of number of events per Myr along each branch based on the time-calibrated phylogeny for *Erebia* (*8*).

To test whether some Merian elements (i.e., ancestral lepidopteran linkage groups; see (*26*)) are more often involved in fusion or fission events than others, we performed chi-square tests to compare observed counts to a uniform distribution. We then tested if some pairs of Merian elements are more likely to fuse together than expected by chance, using a permutation test with 10,000 iterations, keeping the total number of fusions per Merian constant. The *p*-values for each pair were calculated as the proportion of permutations where the permuted counts were equal or higher than the observed counts. *P-*values were then adjusted for multiple testing applying a Benjamini-Hochberg FDR. To test if the number of fusions and fissions in which a Merian element was involved were correlated, we used Spearman’s rank test. We grouped M17 and M20 together in the analyses as they represent an ancient fusion, ancestral to almost all Lepidoptera (*26*). We performed all analyses with R v4.4.1 (*114*) within RStudio v2025.09.2 (*115*).

To test if chromosomal fusions and fissions are associated with repetitive elements expansions, we used Phylogenetic Generalized Least Square (PGLS) regressions between chromosome number, number of fusions and number of fissions along the phylogenetic branches leading to extant species, and genomic proportions of each repeat family. We only considered repeat families for which genomic proportion was > 0.01% in at least 20% of all species (Table S4). We tested the presence of a phylogenetic signal in genomic proportion of each repeat family using the *phylosig* function from the R package *phytools* v2.5-2 (*116*). We performed PGLS regressions using the function *phylolm* from the R package *phylolm* v2.6.5 (*117*) with the lambda model. We also used the PGLS approach to test for general associations between genome size, overall genomic repeat proportion and chromosome number. We then tested if the proportion of each repeat family in the *tyndarus* clade differed from the rest of *Erebia*, by first comparing species of the *tyndarus* clade to all other species and second, comparing the different clades defined in (*8*), using a phylogenetic ANOVA with 10,000 simulations with the function *phylANOVA* from the R package *phytools* v2.4-4 (*116*). For each separate analysis that involved testing all 84 repeat families present in the repeat library, we adjusted for multiple testing applying a false-discovery rate (FDR) correction with the R function *p.adjust*. For the repeat families that were associated with rearrangements or with the *tyndarus* clade, we confirmed the presence of phylogenetic shifts in their genomic proportions using Bayesian reversible-jump multi-optima Ornstein Uhlenbeck models implemented in the R package *bayou* v2.3.0 (*118*) with default priors, 2 million generations and 25% burn-in. Optima shifts were considered significant when their posterior probability was > 0.3.

To test whether the number of rearrangements in which a Merian element is involved is associated with its size or genomic proportion in specific repeat families, we first linked each chromosome in extant species to its corresponding Merian element(s) using the GENESPACE output and the lep_busco_painter tool v.1.0.0 (https://github.com/charlottewright/lep_busco_painter, accessed on 11.12.2025). We then used phylogenetic generalized linear mixed models implemented in the R package *MCMCglmm* v2.36 (*119*), with number of fusions or number of fissions as response variable, Merian size and genomic proportions of each repeat type as explanatory variables, Merian element as a random factor, using a Poisson distribution and taking the phylogeny into account. We only considered repeat families for which genomic proportion was > 0.01% in at least 20% of all species (Table S4). We used 200,000 iterations, a burn-in of 50,000, a thinning interval of 100 and the default priors except for the inverse-Wishart degree of belief parameter (nu = 0.002) and covariance matrix (alpha.V = 1000). We ran four chains in parallel and assessed convergence using the Gelman-Rubin criterion (multivariate potential scale reduction factor < 1.02). As above, we grouped M17 and M20 together.

### Rearrangement breakpoint definition and enrichment tests

We identified rearrangement breakpoints for autosomal fusions and fissions that happened at the tips of the phylogeny and for which we had a closely related species to compare to, resulting in a total of 47 fusion breakpoints (in 11 species pairs) and 64 fission breakpoints (in 3 species pairs). In addition, we considered the 3 fusion breakpoints in the 2 species with heterozygous fusions. We polarized rearrangements into fusions and fissions by inferring the ancestral state from the syngraph output. We defined a breakpoint as the interval between two syntenic regions using the GENESPACE output, in the species with the fused chromosomal state (i.e. ancestral in the case of fission, derived in the case of fusions). We also identified the corresponding homologous regions in the species with the unfused state, often corresponding to subtelomeres defined as the interval between the end of the canonical telomeric repeats and the first synteny block. We identified the canonical lepidopteran telomere repeats “TTAGG” using tidk v0.2.41 (*120*). In some cases where a genomic region was involved in different fusion events in different species, it represented not subtelomeric region but another fusion breakpoint (Fig. 2;Table S11).

To test if rearrangement breakpoints are enriched for specific repetitive elements or segmental duplications, we used GAT v1.3.6 (*121*), which compares the likelihood that repeats overlap with the breakpoint regions compared to 10,000 random genomic regions of similar size in the same chromosome. To make a meaningful comparison between the breakpoints and random genomic regions, we excluded genes using bedtools v2.30.0 (*122*) and the sex chromosomes from the analyses. We tested separately each repeat family for which the genomic proportion was > 0.01% in at least 20% of all species (Table S4) and present both the FDR corrected and uncorrected results. We further tested for an enrichment of repeat families at the two corresponding homologous regions in the species with the unfused state. We considered it as *strong evidence* for an association of a repeat family with a chromosomal rearrangement when the fusion or fission breakpoint and both homologous regions were enriched for the same repeat family (see Fig. 3A). We considered it as *supporting evidence* if either the breakpoint and one homologous region (Fig. 3B) or both homologous regions (Fig. 3C) were enriched for the same repeat family. We tested for enrichment taking either the number of repeats or the number of nucleotides that overlap (i.e. total length of overlaps) into account (Fig. 3D). To be considered strong or supporting evidence, the breakpoint and/or homologous regions all needed to be enriched for the same repeat family with the same criteria. We first tested repeat enrichment separately for each breakpoint. To confirm that some repeat families are repeatedly associated with chromosomal fusions and fissions, we repeated the enrichment analyses grouping all breakpoints in each species pair.

For validation, we repeated the enrichment test using the R package *regioneR* v1.36.0 (*123*) using the function *permTest* with the *circularRandomizeRegions* randomizing function. Contrarily to GAT, *regioneR* compares the enrichment in the region of interest to similar regions genome-wide, rather than per chromosome. We applied the same criteria for strong or supporting evidence as for the GAT analysis. We first analysed each breakpoint separately, then grouped them in a single bed file for each species pair. Enrichment was tested both in terms of number of overlaps, using the *numOverlaps* evaluation function of *regioneR*, and in terms of length of overlaps, using a custom function based on the function *intersect* in the R package *GenomicRanges* 1.56.2 (*124*).

In addition to repetitive elements, we tested if breakpoints and corresponding homologous regions presented the same segmental duplications, defined as genomic regions larger than 1 kb duplicated at least once and having retained at least 90% sequence similarity. We identified segmental duplications using BISER v1.4 (*125*), which has the advantage of simultaneously identifying segmental duplications in species pairs, i.e. if one species presents a segmental duplication, it will look for the same sequence in the other species.

### LINE R1-like sequence and evolution in *Erebia*

To determine if R1-like sequences are longer and more similar between repeat copies at breakpoints compared to 10,000 random genomic regions, we used the function *permTest* with the *resampleRegions* randomizing function and the *meanInRegions* evaluation function in *regioneR*. We ran the comparison once based on median R1-like sequence length and once based on the Kimura 2-parameter (K2P) distance distribution (a proxy for repeat sequence divergence). We calculated K2P for all R1-like elements in each species using the script *divergence_calc.py* from EarlGrey v6.3.3 (*98*). To test for an association between R1-like sequence length or K2P and karyotype, number of fusions, and number of fissions along the phylogeny, we conducted PGLS using the R function *phylolm* with the lambda model. To test for differences between the *tyndarus* clade and other species, and between all different clades, we conducted a phylogenetic ANOVA with 10,000 simulations with the function *phylANOVA* from the R package *phytools* v2.4-4 (*116*).

To investigate which parts of R1-like elements are present at the breakpoints, we first oriented R1-like consensus sequences (5’ – 3’) using either the canonical poly-A tail (defined as at least 9 consecutive As within the first 40 bp at the 3’ end), or another short tandem repeat at the 3’ end detected using trf v4.10 (*126*) with match = 2, mismatch = 7, delta = 7, match probability = 80, indel probability = 10, minimum alignment score = 50 and maximum period = 6. We computed reverse complements of consensus sequences if needed to obtain the 5’ – 3’ orientation using seqkit seq -rp v2.13.0 (*127*). For the R1-like consensus sequences present at breakpoints that could be successfully oriented, we searched for open-reading frames (ORF) using the function getorf from the package EMBOSS v6.6.0.0 (*128*). Two ORFs are expected in complete R1-like sequences (*52*). We confirmed that ORF2 is the expected canonical reverse transcriptase using InterProScan (https://www.ebi.ac.uk/interpro/search/sequence/, accessed on 06.03.2026). To visualise R1-like families, we normalised the coordinates of each R1-like consensus sequence, and their respective ORFs and poly-A or other short tandem repeat tails, from zero to one.

We generated a phylogeny to determine the relatedness among R1-like elements located at breakpoints across *Erebia*. We clustered the consensus sequences for all R1-like elements present at breakpoints to 95% similarity over at least 80% of the length of the shorter sequences to reduce redundancy using mmseqs2 (*129*) easy-cluster (--min-seq-id 0.95 -c 0.8 --cov-mode 1 --cluster-reassign). We then detected ORFs in all six frames in each of these sequences using EMBOSS transeq (-clean -frame 6). We identified matches to known TE domains using an approach described in Baril and Croll (2026). First, we used HMMscan (HMMER v3.4) ‘-E 10 --noal’ to detect homology to known TE protein domains using models supplied in ProfilesBankForREPET_Pfam35.0_GypsyDB, part of the REPET software suite (*131–133*). We filtered hits to retain those with fseq_evalue ≤ 0.001 and fseq_bitscore ≥ 50. We kept only reverse transcriptase (RT) domains by searching for hits with either “RT” in the host protein column or “reverse transcriptase” in the domain description column, or both, and we retained a single hit for each query sequence by selecting the hit with the highest bitscore. This resulted in 98 hits with RT domains. We extracted these sequences from the peptide file generated by EMBOSS transeq, and concatenated them with a FASTA file containing all Pol domains for LINEs (excluding *Penelope*-like elements) from RepeatPeps.lib, which is part of RepeatMasker v4.1.9 (*103*). We aligned the resultant FASTA file with MAFFT v7.526 (*134*) --auto --adjustdirectionaccurately and trimmed with clipkit v2.7.0 (*135*) smart-gap. Finally, we generated a phylogenetic tree of LINE RT domains with iqtree2 v3.0.1 (*136*) with 1000 bootstraps and model Q.pfam+R10, as determined by ModelFinder based on Bayesian information criterion (BIC). We visualized the tree using *ggtree* v3.12.0 (*137*), showing only R1-like elements, and highlighting the sequences present at the breakpoints from other LINE sequences in the dataset.

To visualise the genomic distribution of R1-like elements, we used *karyoploteR* v1.30.0 (*138*) with a window size of 2 Mb. We then tested if R1-like elements, as well as repeats in general, tend to be located closer to the chromosome ends than random genomic regions using *regioneR* with the *randomizeRegions* randomizing function (10,000 randomizations) and the *meanDistance* evaluation function. For species involved in the species pairs analyses, we also tested for repeat enrichment at subtelomeric regions compared to 10,000 random genomic windows using GAT, both in terms of number of overlaps and total length of overlaps.

### Gene family expansions and contractions

To test if gene family expansions and contractions at the base of the *tyndarus* clade could be enriched for functions related to chromosomal rearrangements, we performed a GO enrichment test on gene family expansions and contractions along the *Erebia* phylogeny. We used gene predictions for all 37 *Erebia* genomes (for comparability, we ran BRAKER v3.0.7 for our *Erebia* assemblies without the RNA-seq data, Table S2) and *Maniola jurtina* as outgroup. We used CAFE5 v1.0 (*139*) to identify gene duplications and losses within orthogroups defined by OrthoFinder. We computed a single λ parameter along the phylogeny, removing 16 orthogroups with > 100 copies in at least one species and the orthogroups absent at the root. We used a *p*-value threshold of 0.05 to consider an orthogroup as being expanded or contracted. We then ran topGO v2.56.0 (*140*) to test for an enrichment of Biological Processes GO terms in the sets of genes that were either gained or lost in the *tyndarus* clade compared to the full set of orthogroups. We only considered GO terms with at least five annotated genes and ran topGO with the *weight01* algorithm, which takes the hierarchy of GO terms into account, performing Fisher’s exact test and applying a *p*-value enrichment threshold of 0.01.

To avoid including highly variable gene families, we compared gene family expansions and contractions at the base of the *tyndarus* clade to the ones occurring at the base of the karyotypically conserved *ligea* clade (including *E. eriphyle*, *E. ligea*, *E. euryale*, *E. bubastis* and *E. manto*), and reported only the GO terms enriched in the *tyndarus* but not in the *ligea* gene set.

## Data Availability

All reference genomes and associated raw HiFi, Hi-C and IsoSeq data generated will be deposited at the European Nucleotide Archive under the accession numbers provided in Tables S1 and S2. Gene and repeat annotations, the *Erebia* repeat library, and raw outputs of Genespace, BISER, Fantasia, GAT and regioneR will be deposited on Zenodo. All code is available on GitHub at https://github.com/camille-cornet/Erebia_rearrangements_and_TEs and will be archived on Zenodo. All data will be made available upon acceptance on the manuscript.

## Acknowledgements

We are grateful to Tinatin Chkhartishvili, Giorgi Iankoshvili, Hugo da Costa Vieira, Andreas Sanchez, Martin Albrecht and Pierre Talon for their help in sample collection. We thank the Next Generation Sequencing Platform of the University of Bern for their support in conducting Hi-C and IsoSeq sequencing. We are grateful to the Wellcome Sanger Institute, in particular the Tree of Life Programme, for granting us access to their computational infrastructure to perform the bioinformatic analyses. We further thank the ERGA Community, in particular ERGA-CH, for the useful community resources and guidelines that helped us produce and publish the reference genomes. We thank Catherine Peichel, Daniel Croll, Radu Slobodeanu, Marcial Escudero, Ashwini Mohan, Paula Escuer and Alice Laigle for valuable discussions throughout the preparation of this manuscript.

## Author Contributions

C.C. and K.L. designed the study, acquired funding and collected the samples. C.C., C.W. and K.L. developed the methodology. C.C. and T.B. performed data analyses. C.C., T.B., C.W. and K.L. interpreted the results. K.L. supervised the work. C.C. wrote the original draft, with input from all co-authors. All authors read, revised and approved the final manuscript.

## References

1. E. L. Berdan, T. G. Aubier, S. Cozzolino, R. Faria, J. L. Feder, M. D. Giménez, M. Joron, J. B. Searle, C. Mérot, Structural variants and speciation: multiple processes at play. Cold Spring Harb Perspect Biol 16, a041446 (2024).

2. K. Lucek, M. D. Giménez, M. Joron, M. Rafajlović, J. B. Searle, N. Walden, A. M. Westram, R. Faria, The impact of chromosomal rearrangements in speciation: From micro- to macroevolution. Cold Spring Harb Perspect Biol, a041447 (2023).

3. F. Cicconardi, J. J. Lewis, S. H. Martin, R. D. Reed, C. G. Danko, S. H. Montgomery, Chromosome fusion affects genetic diversity and evolutionary turnover of functional loci but consistently depends on chromosome size. Mol Biol Evol 38, 4449–4462 (2021).

4. K. Näsvall, J. Boman, L. Höök, R. Vila, C. Wiklund, N. Backström, Nascent evolution ofrecombination rate differences as a consequence of chromosomal rearrangements. PLoS Genet 19, e1010717 (2023).

5. R. Faria, A. Navarro, Chromosomal speciation revisited: rearranging theory with pieces of evidence. Trends Ecol Evol 25, 660–669 (2010).

6. Z. Liu, M. Roesti, D. Marques, M. Hiltbrunner, V. Saladin, C. L. Peichel, Chromosomal fusions facilitate adaptation to divergent environments in threespine stickleback. Mol Biol Evol 39, msab358 (2022).

7. Y. Li, S. Wang, Z. Zhang, J. Luo, G. L. Lin, W.-D. Deng, Z. Guo, F. M. Han, L.-L. Wang, J. Li, S.-F. Wu, H.-Q. Liu, S. He, R. W. Murphy, Z.-J. Zhang, D. N. Cooper, D.-D. Wu, Y.-P. Zhang, Large-scale chromosomal changes lead to genome-level expression alterations, environmental adaptation, and speciation in the Gayal (*Bos frontalis*). Mol Biol Evol 40, msad006 (2023).

8. H. Augustijnen, L. Bätscher, M. Cesanek, T. Chkhartishvili, V. Dincă, G. Iankoshvili, K. Ogawa, R. Vila, S. Klopfstein, J. M. de Vos, K. Lucek, A macroevolutionary role for chromosomal fusion and fission in *Erebia* butterflies. Sci Adv 10, eadl0989 (2024).

9. A. Carta, M. Escudero, Karyotypic diversity: a neglected trait to explain angiosperm diversification? Evol 77, 1158–1164 (2023).

10. E. Olmo, Rate of chromosome changes and speciation in reptiles. Genetica 125, 185–203 (2005).

11. T. D. Lewin, I. J.-Y. Liao, Y.-J. Luo, Conservation of bilaterian genome structure is the exception, not the rule. Genome Biol 26, 247 (2025).

12. O. Simakov, J. Bredeson, K. Berkoff, F. Marletaz, T. Mitros, D. T. Schultz, B. L. O’Connell, P. Dear, D. E. Martinez, R. E. Steele, R. E. Green, C. N. David, D. S. Rokhsar, Deeply conserved synteny and the evolution of metazoan chromosomes. Sci Adv 8, eabi5884 (2022).

13. M. Muffato, A. Louis, N. T. T. Nguyen, J. Lucas, C. Berthelot, H. Roest Crollius, Reconstruction of hundreds of reference ancestral genomes across the eukaryotic kingdom. Nat Ecol Evol 7, 355–366 (2023).

14. T. D. Lewin, I. J.-Y. Liao, Y.-J. Luo, Annelid comparative genomics and the evolution of massive lineage-specific genome rearrangement in bilaterians. Mol Biol Evol 41, msae172 (2024).

15. D. T. Schultz, E. A. C. Heath-Heckman, C. J. Winchell, D.-H. T. Kuo, Y. Yu, F. Oberauer, K. Kocot, S.-J. Cho, O. Simakov, D. A. Weisblat, Acceleration of genome rearrangement in clitellate annelids. bioRxiv [Preprint] (2024). 10.1101/2024.05.12.593736.

16. C. Vargas-Chávez, L. Benítez-Álvarez, G. I. Martínez-Redondo, L. Álvarez-González, J. Salces-Ortiz, K. Eleftheriadi, N. Escudero, N. Guiglielmoni, J.-F. Flot, M. Novo, A. Ruiz-Herrera, A. McLysaght, R. Fernández, An episodic burst of massive genomic rearrangements and the origin of non-marine annelids. Nat Ecol Evol 9, 1263–1279 (2025).

17. S. Garagna, J. Page, R. Fernandez-Donoso, M. Zuccotti, J. B. Searle, The Robertsonian phenomenon in the house mouse: mutation, meiosis and speciation. Chromosoma 123, 529–544 (2014).

18. J. L. Gerton, A working model for the formation of Robertsonian chromosomes. J Cell Sci 137, jcs261912 (2024).

19. A. Marques, I. A. Drinnenberg, Same but different: Centromere regulations in holocentric insects and plants. Curr Opin Cell Biol 93, 102484 (2025).

20. D. P. Melters, L. V. Paliulis, I. F. Korf, S. W. L. Chan, Holocentric chromosomes: convergent evolution, meiotic adaptations, and genomic analysis. Chromosome Res 20, 579–593 (2012).

21. K. Lucek, H. Augustijnen, M. Escudero, A holocentric twist to chromosomal speciation? Trends Ecol Evol 37, 655–662 (2022).

22. V. A. Lukhtanov, V. Dincă, M. Friberg, J. Šíchová, M. Olofsson, R. Vila, F. Marec, C. Wiklund, Versatility of multivalent orientation, inverted meiosis, and rescued fitness in holocentric chromosomal hybrids. Proc Natl Acad Sci 115, E9610–E9619 (2018).

23. M. Mandrioli, G. C. Manicardi, Holocentric chromosomes. PLoS Genet 16, e1008918 (2020).

24. A. Y. Kawahara, D. Plotkin, M. Espeland, K. Meusemann, E. F. A. Toussaint, A. Donath, F. Gimnich, P. B. Frandsen, A. Zwick, M. dos Reis, J. R. Barber, R. S. Peters, S. Liu, X. Zhou, C. Mayer, L. Podsiadlowski, C. Storer, J. E. Yack, B. Misof, J. W. Breinholt, Phylogenomics reveals the evolutionary timing and pattern of butterflies and moths. Proc Natl Acad Sci 116, 22657–22663 (2019).

25. C. J. Wright, V. M. Shirey, F. L. Condamine, J. K. Hill, N. E. Pierce, N. Wahlberg, A. Y. Kawahara, Evolution, genomics and conservation of butterflies and moths. Nat Rev Biodivers, 1–18 (2026).

26. C. J. Wright, L. Stevens, A. Mackintosh, M. Lawniczak, M. Blaxter, Comparative genomics reveals the dynamics of chromosome evolution in Lepidoptera. Nat Ecol Evol, 1–14 (2024).

27. J. M. de Vos, H. Augustijnen, L. Bätscher, K. Lucek, Speciation through chromosomal fusion and fission in Lepidoptera. Philos Trans R Soc B 375, 20190539 (2020).

28. C. Peña, H. Witthauer, I. Klečková, Z. Fric, N. Wahlberg, Adaptive radiations in butterflies: evolutionary history of the genus *Erebia* (Nymphalidae: Satyrinae). Biol J Linn Soc 116, 449–467 (2015).

29. I. Klečková, J. Klečka, Z. F. Fric, M. Česánek, L. Dutoit, L. Pellissier, P. Matos-Maraví, Climatic niche conservatism and ecological diversification in the holarctic cold-dwelling butterfly genus *Erebia*. Insect Syst Divers 7, 2 (2023).

30. K. Lucek, Evolutionary mechanisms of varying chromosome numbers in the radiation of *Erebia* butterflies. Genes 9, 166 (2018).

31. K. Lucek, D. Linke, C. Wright, J. Meier, M. Blaxter, The genome sequence of the Styrian Ringlet, *Erebia stirius* Godart, 1823 (Lepidoptera: Nymphalidae) [version 1; peer review: 1 approved with reservations]. Wellcome Open Research 10 (2025).

32. C. Diblasi, M. Saitou, Beyond inversions and deletions: the evolutionary and functional insights from translocations, fissions, and fusions in animal genomes. Heredity, 1–12 (2025).

33. W.-E. Lönnig, H. Saedler, Chromosome rearrangements and transposable elements. Annu Rev Genet 36, 389–410 (2002).

34. M. Cáceres, J. M. Ranz, A. Barbadilla, M. Long, A. Ruiz, Generation of a widespread *Drosophila* inversion by a transposable element. Science 285, 415–418 (1999).

35. A. Delprat, B. Negre, M. Puig, A. Ruiz, The transposon Galileo generates natural chromosomal inversions in *Drosophila* by ectopic recombination. PLOS ONE 4, e7883 (2009).

36. P. Balachandran, I. A. Walawalkar, J. I. Flores, J. N. Dayton, P. A. Audano, C. R. Beck, Transposable element-mediated rearrangements are prevalent in human genomes. Nat Commun 13, 7115 (2022).

37. R. Aasegg Araya, W. B. Reinar, O. K. Tørresen, C. Goubert, T. J. Daughton, S. N. K. Hoff, H. T. Baalsrud, M. S. O. Brieuc, A. Z. Komisarczuk, S. Jentoft, J. Cerca, K. S. Jakobsen, Chromosomal inversions mediated by tandem insertions of transposable elements. Genome Biol Evol 17, evaf131 (2025).

38. L. Gozashti, O. S. Harringmeyer, H. E. Hoekstra, How repeats rearrange chromosomes: The molecular basis of chromosomal inversions in deer mice. Cell Reports 44, 115644 (2025).

39. L. G. de Lima, A. Guarracino, S. Koren, T. Potapova, S. McKinney, A. Rhie, S. J. Solar, C. Seidel, B. L. Fagen, B. P. Walenz, G. G. Bouffard, S. Y. Brooks, M. Peterson, K. Hall, J. Crawford, A. C. Young, B. D. Pickett, E. Garrison, A. M. Phillippy, J. L. Gerton, The formation and propagation of human Robertsonian chromosomes. Nature 647, 952–961 (2025).

40. P. G. Hofstatter, G. Thangavel, T. Lux, P. Neumann, T. Vondrak, P. Novak, M. Zhang, L. Costa, M. Castellani, A. Scott, H. Toegelová, J. Fuchs, Y. Mata-Sucre, Y. Dias, A. L. L. Vanzela, B. Huettel, C. C. S. Almeida, H. Šimková, G. Souza, A. Pedrosa-Harand, J. Macas, K. F. X. Mayer, A. Houben, A. Marques, Repeat-based holocentromeres influence genome architecture and karyotype evolution. Cell 185, 3153–3168.e18 (2022).

41. J. I. McCulloch, M. Uliano-Silva, C. J. Wright, I. R. Henderson, S. Ebdon, D. T. of L. Consortium, K. S. Jaron, M. Blaxter, Rapid chromosomal evolution and oligocentromeric drive in sedges and rushes. bioRxiv [Preprint] (2026). 10.64898/2026.02.27.708619.

42. A. P. Senaratne, H. Muller, K. A. Fryer, M. Kawamoto, S. Katsuma, I. A. Drinnenberg, Formation of the CenH3-deficient holocentromere in Lepidoptera avoids active chromatin. Curr Biol 31, 173–181.e7 (2021).

43. C. J. Wright, N. Wahlberg, R. Vila, M. Mutanen, P. Matos-Maraví, K. Lucek, I. Kleckova, L. Dapporto, V. Dincă, C. Bruschini, C. W. Wheat, M. Vila, L. Torrado-Blanco, V. Todisco, M. Rindos, P. Nguyen, P. O. Mulhair, S. Kamenova, M. Hicks, M. Espeland, I. A. Drinnenberg, M. Doblas-Bajo, R. I. Bailey, M. Blaxter, J. I. Meier, G. Zucco, M. Zlatkovic, M. Wiorek, M. Wiemers, C. C. Weber, P. Vrba, V. Visacki, J. V. P. S. Rita, M. Vallinmäki, B. Ulaşlı, E. Toro-Delgado, G. G. Thallinger, A. S. Bartonova, J. Stewart, C. Spilker, L. Spilani, D. Schmitz, B. Schattanek-Wiesmar, P. Schattanek-Wiesmair, S. Scalercio, G. S. Sramkó, D. V. Stojanovic, C. G. Sotero-Caio, J. Rota, M. Repullés, Ł. Przybyłowicz, P. Potocký, J. Pohjoismäki, E. Õunap, A. S. Ortiz, R. Nunez, V. Nazari, A. Nahirnić-Beshkova, T. Moreira, M. Meyer, M. Menchetti, D. McKenna, V. Marques, K. Lohse, L. Littmann, D. Linke, J. Lancho, S. L. Cava, E. Kriegova, E. Keskin, S. Kamenova, M. Joron, A. K. Hundsdoerfer, N. Hévin, J. Gauthier, M. Fabusova, Z. F. Fric, P. Escuer, B. Emerson, M. Elias, L. Downes, M. Doblas-Bajo, L. Despres, T. Decroly, C. Cornet, T. Chkhartishvili, R. Challis, J. Bridle, B. Boussau, A. Bordoni, J. Boman, S. Beshkov, F. F. Bertocchini, S. Baral, J. Baixeras, D. Absolon, Project Psyche: reference genomes for all Lepidoptera in Europe. Trends Ecol Evol 40, 1234–1250 (2025).

44. A. Rhie, S. A. McCarthy, O. Fedrigo, J. Damas, G. Formenti, S. Koren, M. Uliano-Silva, W. Chow, A. Fungtammasan, J. Kim, C. Lee, B. J. Ko, M. Chaisson, G. L. Gedman, L. J. Cantin, F. Thibaud-Nissen, L. Haggerty, I. Bista, M. Smith, B. Haase, J. Mountcastle, S. Winkler, S. Paez, J. Howard, S. C. Vernes, T. M. Lama, F. Grutzner, W. C. Warren, C. N. Balakrishnan, D. Burt, J. M. George, M. T. Biegler, D. Iorns, A. Digby, D. Eason, B. Robertson, T. Edwards, M. Wilkinson, G. Turner, A. Meyer, A. F. Kautt, P. Franchini, H. W. Detrich, H. Svardal, M. Wagner, G. J. P. Naylor, M. Pippel, M. Malinsky, M. Mooney, M. Simbirsky, B. T. Hannigan, T. Pesout, M. Houck, A. Misuraca, S. B. Kingan, R. Hall, Z. Kronenberg, I. Sović, C. Dunn, Z. Ning, A. Hastie, J. Lee, S. Selvaraj, R. E. Green, N. H. Putnam, I. Gut, J. Ghurye, E. Garrison, Y. Sims, J. Collins, S. Pelan, J. Torrance, A. Tracey, J. Wood, R. E. Dagnew, D. Guan, S. E. London, D. F. Clayton, C. V. Mello, S. R. Friedrich, P. V. Lovell, E. Osipova, F. O. Al-Ajli, S. Secomandi, H. Kim, C. Theofanopoulou, M. Hiller, Y. Zhou, R. S. Harris, K. D. Makova, P. Medvedev, J. Hoffman, P. Masterson, K. Clark, F. Martin, K. Howe, P. Flicek, B. P. Walenz, W. Kwak, H. Clawson, M. Diekhans, L. Nassar, B. Paten, R. H. S. Kraus, A. J. Crawford, M. T. P. Gilbert, G. Zhang, B. Venkatesh, R. W. Murphy, K.-P. Koepfli, B. Shapiro, W. E. Johnson, F. Di Palma, T. Marques-Bonet, E. C. Teeling, T. Warnow, J. M. Graves, O. A. Ryder, D. Haussler, S. J. O’Brien, J. Korlach, H. A. Lewin, K. Howe, E. W. Myers, R. Durbin, A. M. Phillippy, E. D. Jarvis, Towards complete and error-free genome assemblies of all vertebrate species. Nature 592, 737–746 (2021).

45. D. A. S. Smith, I. J. Gordon, W. Traut, J. Herren, S. Collins, D. J. Martins, K. Saitoti, P. Ireri, R. ffrench-Constant, A neo-W chromosome in a tropical butterfly links colour pattern, male-killing, and speciation. Proc R Soc B 283, 20160821 (2016).

46. H. An, K. Nam, Lepidopteran genomes have denser transposable elements in smaller chromosomes, likely driven by non-allelic homologous recombination. Genome Biol Evol 17, evaf137 (2025).

47. F. Thörn, J.-L. Claret, N. Backström, J. Boman, Determinants of chromosomal rearrangements in holocentric *Leptidea* butterflies. bioRxiv [Preprint] (2026). 10.64898/2026.02.26.708211.

48. Z. Kuang, X. Yang, N. Wan, J. Chen, Q. Duan, B. Li, X. Liu, X. Liang, X. Liu, W. Liu, E. Nevo, K. Li, Genomic insights into chromosomal fusion and its evolutionary implications for zokors. Mol Biol Evol 43, msag032 (2026).

49. L. Lopez-Delisle, L. Rabbani, J. Wolff, V. Bhardwaj, R. Backofen, B. Grüning, F. Ramírez, T. Manke, pyGenomeTracks: reproducible plots for multivariate genomic datasets. Bioinformatics 37, 422–423 (2021).

50. J. N. Wells, C. Feschotte, A field guide to eukaryotic transposable elements. Annu Rev Genet 54, 539–561 (2020).

51. J. Renkawitz, C. A. Lademann, S. Jentsch, Mechanisms and principles of homology search during recombination. Nat Rev Mol Cell Biol 15, 369–383 (2014).

52. C. E. Pérez-González, T. H. Eickbush, Dynamics of R1 and R2 elements in the rDNA locus of *Drosophila simulans*. Genetics 158, 1557–1567 (2001).

53. V. A. Lukhtanov, E. A. Pazhenkova, Diversity and evolution of telomeric motifs and telomere DNA organization in insects. Biol J Linn Soc 140, 536–555 (2023).

54. C. Cornet, P. Mora, H. Augustijnen, P. Nguyen, M. Escudero, K. Lucek, Holocentric repeat landscapes: From micro-evolutionary patterns to macro-evolutionary associations with karyotype evolution. Mol Ecol 33, e17100 (2024).

55. L. Hilgers, M. Rovatsos, D.-G. Kontopoulos, T. Brown, T. Hickler, B. Huntley, M. Pippel, C. Munegowda, T. Mueller, A. Ahmed, A. Laas, P. Praschag, J. Damas, S. Winkler, H. Lewin, E. Myers, U. Fritz, M. Hiller, Side-necked turtle genomes reveal chromosomal dynamics, skeletal innovation and cancer resistance. bioRxiv [Preprint] (2026). 10.64898/2026.03.05.709825.

56. T. C. Mathers, R. H. M. Wouters, S. T. Mugford, D. Swarbreck, C. van Oosterhout, S. A. Hogenhout, Chromosome-scale genome assemblies of aphids reveal extensively rearranged autosomes and long-term conservation of the X chromosome. Mol Biol Evol 38, 856–875 (2021).

57. L. Höök, K. Näsvall, R. Vila, C. Wiklund, N. Backström, High-density linkage maps and chromosome level genome assemblies unveil direction and frequency of extensive structural rearrangements in wood white butterflies (*Leptidea* spp.). Chromosome Res 31, 2 (2023).

58. C. J. Wright, D. Absolon, M. Gascoigne-Pees, R. Vila, M. K. N. Lawniczak, M. Blaxter, Constraints on chromosome evolution revealed by the 229 chromosome pairs of the Atlas blue butterfly. Curr Biol 35, 4727–4742.e7 (2025).

59. V. Ahola, R. Lehtonen, P. Somervuo, L. Salmela, P. Koskinen, P. Rastas, N. Välimäki, L. Paulin, J. Kvist, N. Wahlberg, J. Tanskanen, E. A. Hornett, L. C. Ferguson, S. Luo, Z. Cao, M. A. de Jong, A. Duplouy, O.-P. Smolander, H. Vogel, R. C. McCoy, K. Qian, W. S. Chong, Q. Zhang, F. Ahmad, J. K. Haukka, A. Joshi, J. Salojärvi, C. W. Wheat, E. Grosse-Wilde, D. Hughes, R. Katainen, E. Pitkänen, J. Ylinen, R. M. Waterhouse, M. Turunen, A. Vähärautio, S. P. Ojanen, A. H. Schulman, M. Taipale, D. Lawson, E. Ukkonen, V. Mäkinen, M. R. Goldsmith, L. Holm, P. Auvinen, M. J. Frilander, I. Hanski, The Glanville fritillary genome retains an ancient karyotype and reveals selective chromosomal fusions in Lepidoptera. Nat Commun 5, 4737 (2014).

60. H. Augustijnen, T. Patsiou, K. Lucek, Secondary contact rather than coexistence—*Erebia* butterflies in the Alps. Evolution 76, 2669–2686 (2022).

61. H. Augustijnen, K. Lucek, Beyond gene flow: (non)-parallelism of secondary contact in a pair of highly differentiated sibling species. Mol Ecol 33, e17488 (2024).

62. E. S. M. van der Heijden, K. Näsvall, F. A. Seixas, C. E. Beserra Nobre, A. C. D. Maia, P. Salazar-Carrión, J. M. Walker, D. Szczerbowski, S. Schulz, I. A. Warren, K. G. Gavilanes Córdova, M. J. Sánchez-Carvajal, F. Chandi, A. P. Arias-Cruz, N. Rueda-M, C. Salazar, K. K. Dasmahapatra, S. H. Montgomery, M. McClure, D. E. Absolon, T. C. Mathers, C. A. Santos, S. McCarthy, J. M. D. Wood, G. Lamas, C. Bacquet, A. V. L. Freitas, K. R. Willmott, C. D. Jiggins, M. Elias, J. I. Meier, Genomics of Neotropical biodiversity indicators: Two butterfly radiations with rampant chromosomal rearrangements and hybridization. Proc Natl Acad Sci 122, e2410939122 (2025).

63. C. M. Tribble, J. I. Márquez-Corro, M. R. May, A. L. Hipp, M. Escudero, R. Zenil-Ferguson, Macroevolutionary inference of complex modes of chromosomal speciation in a cosmopolitan plant lineage. New Phytologist 245, 2350–2361 (2025).

64. J. Damas, M. Corbo, J. Kim, J. Turner-Maier, M. Farré, D. M. Larkin, O. A. Ryder, C. Steiner, M. L. Houck, S. Hall, L. Shiue, S. Thomas, T. Swale, M. Daly, J. Korlach, M. Uliano-Silva, C. J. Mazzoni, B. W. Birren, D. P. Genereux, J. Johnson, K. Lindblad-Toh, E. K. Karlsson, M. T. Nweeia, R. N. Johnson, Zoonomia Consortium, H. A. Lewin, Evolution of the ancestral mammalian karyotype and syntenic regions. Proc Natl Acad Sci 119, e2209139119 (2022).

65. A. Mackintosh, R. Vila, S. H. Martin, D. Setter, K. Lohse, Do chromosome rearrangements fix by genetic drift or natural selection? Insights from *Brenthis* butterflies. Mol Ecol 33, e17146 (2024).

66. D. C. Presgraves, Evaluating genomic signatures of “the large X-effect” during complex speciation. Mol Ecol 27, 3822–3830 (2018).

67. A. K. Huylmans, A. Macon, B. Vicoso, Global dosage compensation is ubiquitous in Lepidoptera, but counteracted by the masculinization of the Z chromosome. Mol Biol Evol 34, 2637–2649 (2017).

68. W. Traut, K. Sahara, F. Marec, Sex chromosomes and sex determination in Lepidoptera. Sex Dev 1, 332–346 (2007).

69. G. Pascarella, C. C. Hon, K. Hashimoto, A. Busch, J. Luginbühl, C. Parr, W. H. Yip, K. Abe, A. Kratz, A. Bonetti, F. Agostini, J. Severin, S. Murayama, Y. Suzuki, S. Gustincich, M. Frith, P. Carninci, Recombination of repeat elements generates somatic complexity in human genomes. Cell 185, 3025–3040.e6 (2022).

70. C. Robberecht, T. Voet, M. Z. Esteki, B. A. Nowakowska, J. R. Vermeesch, Nonallelic homologous recombination between retrotransposable elements is a driver of de novo unbalanced translocations. Genome Res. 23, 411–418 (2013).

71. M. Startek, P. Szafranski, T. Gambin, I. M. Campbell, P. Hixson, C. A. Shaw, P. Stankiewicz, A. Gambin, Genome-wide analyses of LINE–LINE-mediated nonallelic homologous recombination. Nucleic Acids Res 43, 2188–2198 (2015).

72. J. S. Sproul, S. Hotaling, J. Heckenhauer, A. Powell, D. Marshall, A. M. Larracuente, J. L. Kelley, S. U. Pauls, P. B. Frandsen, Analyses of 600+ insect genomes reveal repetitive element dynamics and highlight biodiversity-scale repeat annotation challenges. Genome Res 33, 1708–1717 (2023).

73. K. K. Kojima, H. Fujiwara, Evolution of target specificity in R1 clade non-LTR retrotransposons. Mol Biol Evol 20, 351–361 (2003).

74. H. Fujiwara, M. Osanai, T. Matsumoto, K. K. Kojima, Telomere-specific non-LTR retrotransposons and telomere maintenance in the silkworm, *Bombyx mori*. Chromosome Res 13, 455–467 (2005).

75. A. I. Kalmykova, O. A. Sokolova, Retrotransposons and telomeres. Biochemistry Moscow 88, 1739–1753 (2023).

76. T. Wicker, F. Sabot, A. Hua-Van, J. L. Bennetzen, P. Capy, B. Chalhoub, A. Flavell, P. Leroy, M. Morgante, O. Panaud, E. Paux, P. SanMiguel, A. H. Schulman, A unified classification system for eukaryotic transposable elements. Nat Rev Genet 8, 973–982 (2007).

77. D. A. Petrov, Y. T. Aminetzach, J. C. Davis, D. Bensasson, A. E. Hirsh, Size matters: Non-LTR retrotransposable elements and ectopic recombination in *Drosophila*. Mol Biol Evol 20, 880–892 (2003).

78. M. S. Longo, D. M. Carone, E. D. Green, M. J. O’Neill, R. J. O’Neill, NISC Comparative Sequencing Program, Distinct retroelement classes define evolutionary breakpoints demarcating sites of evolutionary novelty. BMC Genomics 10, 334 (2009).

79. X. Yu, A. Gabriel, Reciprocal translocations in *Saccharomyces cerevisiae* formed by nonhomologous end joining. Genetics 166, 741–751 (2004).

80. B. Burssed, M. Zamariolli, F. T. Bellucco, M. I. Melaragno, Mechanisms of structural chromosomal rearrangement formation. Mol Cytogenet 15, 23 (2022).

81. S. Zumalave, M. Santamarina, N. P. Espasandín, J. Zamora, D. Garcia-Souto, J. Temes, T. M. Baker, J. Rodríguez-Castro, P. Otero, A. Pequeño-Valtierra, I. Otero, A. Oitabén, E. G. Álvarez, I. Díaz-Arias, M. Martínez-Fernández, M. G. Blanco, P. Van Loo, G. Cristofari, B. Rodriguez-Martin, J. M. C. Tubio, Concurrent L1 retrotransposition events promote reciprocal translocations in human tumorigenesis. Science 392, eaee4513 (2026).

82. S. K. Bohlander, P. M. Kakadia, “DNA repair and chromosomal translocations” in Chromosomal Instability in Cancer Cells, B. M. Ghadimi, T. Ried, Eds. (Springer International Publishing Switzerland, Cham, 2015; 10.1007/978-3-319-20291-4_1), pp. 1–37.

83. K. Zhao, S. Y. Rhee, Interpreting omics data with pathway enrichment analysis. Trends Genet 39, 308–319 (2023).

84. Z. Huang, I. De O. Furo, J. Liu, V. Peona, A. J. B. Gomes, W. Cen, H. Huang, Y. Zhang, D. Chen, T. Xue, Q. Zhang, Z. Yue, Q. Wang, L. Yu, Y. Chen, A. Suh, E. H. C. de Oliveira, L. Xu, Recurrent chromosome reshufling and the evolution of neo-sex chromosomes in parrots. Nat Commun 13, 944 (2022).

85. N. Wan, Q. Duan, Z. Cai, Z. Zhu, J. Wang, Y. Tian, W. Shen, B. Li, Z. Kuang, X. Liang, S. Liu, X. An, X. Yang, X. Liu, L. Mao, J. Chen, Y. Wang, Z. Feng, W. Liu, Y. Bu, E. Nevo, R. Papa, A. Meyer, J. Liu, K. Li, Aplf/Dna2 variants drive chromosomal fission and accelerate speciation in zokors. Sci Adv 11, eadt2282 (2025).

86. L. Carbone, R. Alan Harris, S. Gnerre, K. R. Veeramah, B. Lorente-Galdos, J. Huddleston, T. J. Meyer, J. Herrero, C. Roos, B. Aken, F. Anaclerio, N. Archidiacono, C. Baker, D. Barrell, M. A. Batzer, K. Beal, A. Blancher, C. L. Bohrson, M. Brameier, M. S. Campbell, O. Capozzi, C. Casola, G. Chiatante, A. Cree, A. Damert, P. J. de Jong, L. Dumas, M. Fernandez-Callejo, P. Flicek, N. V. Fuchs, I. Gut, M. Gut, M. W. Hahn, J. Hernandez-Rodriguez, L. W. Hillier, R. Hubley, B. Ianc, Z. Izsvák, N. G. Jablonski, L. M. Johnstone, A. Karimpour-Fard, M. K. Konkel, D. Kostka, N. H. Lazar, S. L. Lee, L. R. Lewis, Y. Liu, D. P. Locke, S. Mallick, F. L. Mendez, M. Muffato, L. V. Nazareth, K. A. Nevonen, M. O’Bleness, C. Ochis, D. T. Odom, K. S. Pollard, J. Quilez, D. Reich, M. Rocchi, G. G. Schumann, S. Searle, J. M. Sikela, G. Skollar, A. Smit, K. Sonmez, B. ten Hallers, E. Terhune, G. W. C. Thomas, B. Ullmer, M. Ventura, J. A. Walker, J. D. Wall, L. Walter, M. C. Ward, S. J. Wheelan, C. W. Whelan, S. White, L. J. Wilhelm, A. E. Woerner, M. Yandell, B. Zhu, M. F. Hammer, T. Marques-Bonet, E. E. Eichler, L. Fulton, C. Fronick, D. M. Muzny, W. C. Warren, K. C. Worley, J. Rogers, R. K. Wilson, R. A. Gibbs, Gibbon genome and the fast karyotype evolution of small apes. Nature 513, 195–201 (2014).

87. M. Schartl, J. M. Woltering, I. Irisarri, K. Du, S. Kneitz, M. Pippel, T. Brown, P. Franchini, J. Li, M. Li, M. Adolfi, S. Winkler, J. de Freitas Sousa, Z. Chen, S. Jacinto, E. Z. Kvon, L. R. Correa de Oliveira, E. Monteiro, D. Baia Amaral, T. Burmester, D. Chalopin, A. Suh, E. Myers, O. Simakov, I. Schneider, A. Meyer, The genomes of all lungfish inform on genome expansion and tetrapod evolution. Nature 634, 96–103 (2024).

88. A. V. Mohan, P. Escuer, C. Cornet, K. Lucek, A three-dimensional genomics view for speciation research. Trends Genet 40, 638–641 (2024).

89. S. Bouaouina, Y. Chittaro, Y. Willi, K. Lucek, Asynchronous life cycles contribute to reproductive isolation between two Alpine butterflies. Evolution Letters, qrad046 (2023).

90. Darwin Tree of Life Project Consortium, Sequence locally, think globally: The Darwin Tree of Life Project. Proc Natl Acad Sci 119, e2115642118 (2022).

91. M. Vila, L. Torrado-Blanco, N. Ryrholm, D. Romero-Pedreira, M. Conejero, A. Böhne, R. Monteiro, T. Marcussen, R. A. Oomen, T. H. Struck, L. Aguilera, M. Gut, F. Câmara Ferreira, F. Cruz, J. Gómez-Garrido, T. S. Alioto, L. Haggerty, F. Martin, C. Bortoluzzi, ERGA-BGE genome of *Erebia palarica* Chapman, 1905: a montane butterfly endemic to North-West Spain. Open Res Europe 6, 9 (2026).

92. C. Cornet, K. Lucek, The genome sequence of the Common Brassy Ringlet, Erebia cassioides (Reiner & Hohenwarth, 1792) (Lepidoptera, Nymphalidae). Res Ideas Outcomes 11, e174988 (2025).

93. H. Cheng, G. T. Concepcion, X. Feng, H. Zhang, H. Li, Haplotype-resolved de novo assembly using phased assembly graphs with hifiasm. Nat Methods 18, 170–175 (2021).

94. D. Guan, S. A. McCarthy, J. Wood, K. Howe, Y. Wang, R. Durbin, Identifying and removing haplotypic duplication in primary genome assemblies. Bioinformatics 36, 2896–2898 (2020).

95. C. Zhou, S. A. McCarthy, R. Durbin, YaHS: yet another Hi-C scaffolding tool. Bioinformatics 39, btac808 (2023).

96. N. C. Durand, J. T. Robinson, M. S. Shamim, I. Machol, J. P. Mesirov, E. S. Lander, E. L. Aiden, Juicebox provides a visualization system for Hi-C contact maps with unlimited zoom. Cell Syst 3, 99–101 (2016).

97. M. Uliano-Silva, J. G. R. N. Ferreira, K. Krasheninnikova, M. Blaxter, N. Mieszkowska, N. Hall, P. Holland, R. Durbin, T. Richards, P. Kersey, P. Hollingsworth, W. Wilson, A. Twyford, E. Gaya, M. Lawniczak, O. Lewis, G. Broad, F. Martin, M. Hart, I. Barnes, G. Formenti, L. Abueg, J. Torrance, E. W. Myers, R. Durbin, M. Blaxter, S. A. McCarthy, Darwin Tree of Life Consortium, MitoHiFi: a python pipeline for mitochondrial genome assembly from PacBio high fidelity reads. BMC Bioinform 24, 288 (2023).

98. T. Baril, J. Galbraith, A. Hayward, Earl Grey: A fully automated user-friendly transposable element annotation and analysis pipeline. Mol Biol Evol 41, msae068 (2024).

99. J. Storer, R. Hubley, J. Rosen, T. J. Wheeler, A. F. Smit, The Dfam community resource of transposable element families, sequence models, and genome annotations. Mobile DNA 12, 2 (2021).

100. W. Li, A. Godzik, Cd-hit: a fast program for clustering and comparing large sets of protein or nucleotide sequences. Bioinformatics 22, 1658–1659 (2006).

101. C. Goubert, R. J. Craig, A. F. Bilat, V. Peona, A. A. Vogan, A. V. Protasio, A beginner’s guide to manual curation of transposable elements. Mobile DNA 13, 7 (2022).

102. B. Buchfink, K. Reuter, H.-G. Drost, Sensitive protein alignments at tree-of-life scale using DIAMOND. Nat Methods 18, 366–368 (2021).

103. A. Smit, R. Hubley, P. Green, RepeatMasker Open-4.0., (2013); http://www.repeatmasker.org.

104. W. De Coster, R. Rademakers, NanoPack2: population-scale evaluation of long-read sequencing data. Bioinformatics 39, btad311 (2023).

105. H. Li, Minimap2: pairwise alignment for nucleotide sequences. Bioinformatics 34, 3094–3100 (2018).

106. R. M. Waterhouse, F. Tegenfeldt, J. Li, E. M. Zdobnov, E. V. Kriventseva, OrthoDB: a hierarchical catalog of animal, fungal and bacterial orthologs. Nucleic Acids Res 41, D358–D365 (2013).

107. T. Brůna, L. Gabriel, K. J. Hoff, Navigating eukaryotic genome annotation pipelines: A route map to BRAKER, Galba, and TSEBRA. arXiv arXiv:2403.19416 [Preprint] (2024). 10.48550/arXiv.2403.19416.

108. T. Brůna, K. J. Hoff, A. Lomsadze, M. Stanke, M. Borodovsky, BRAKER2: automatic eukaryotic genome annotation with GeneMark-EP+ and AUGUSTUS supported by a protein database. NAR Genom Bioinform 3, lqaa108 (2021).

109. M. Manni, M. R. Berkeley, M. Seppey, F. A. Simão, E. M. Zdobnov, BUSCO update: Novel and streamlined workflows along with broader and deeper phylogenetic coverage for scoring of eukaryotic, prokaryotic, and viral genomes. Mol Biol Evol 38, 4647–4654 (2021).

110. G. I. Martínez-Redondo, F. M. Perez-Canales, B. Carbonetto, J. M. Fernández, I. Barrios-Núñez, M. Vázquez-Valls, I. Cases, A. M. Rojas, R. Fernández, FANTASIA leverages language models to decode the functional dark proteome across the animal tree of life. Commun Biol 8, 1–8 (2025).

111. D. M. Emms, S. Kelly, OrthoFinder: phylogenetic orthology inference for comparative genomics. Genome Biol 20, 238 (2019).

112. J. T. Lovell, A. Sreedasyam, M. E. Schranz, M. Wilson, J. W. Carlson, A. Harkess, D. Emms, D. M. Goodstein, J. Schmutz, GENESPACE tracks regions of interest and gene copy number variation across multiple genomes. eLife 11, e78526 (2022).

113. A. Mackintosh, P. M. G. de la Rosa, S. H. Martin, K. Lohse, D. R. Laetsch, Inferring inter-chromosomal rearrangements and ancestral linkage groups from synteny. bioRxiv [Preprint] (2023). 10.1101/2023.09.17.558111.

114. R Core Team, R: A Language and Environment for Statistical Computing (R Foundation for Statistical Computing, Vienna, Austria, 2022; https://www.R-project.org/).

115. Posit team, “RStudio: Integrated development environment for R” (manual, Boston, MA, 2023); http://www.posit.co/.

116. L. J. Revell, phytools 2.0: an updated R ecosystem for phylogenetic comparative methods (and other things). PeerJ 12, e16505 (2024).

117. L. si T. Ho, C. Ané, A linear-time algorithm for gaussian and non-gaussian trait evolution models. Syst Biol 63, 397–408 (2014).

118. J. C. Uyeda, L. J. Harmon, A novel bayesian method for inferring and interpreting the dynamics of adaptive landscapes from phylogenetic comparative data. Syst Biol 63, 902–918 (2014).

119. J. D. Hadfield, MCMC methods for multi-response generalized linear mixed models: The MCMCglmm R package. J Stat Softw 33, 1–22 (2010).

120. M. R. Brown, P. Manuel Gonzalez de La Rosa, M. Blaxter, tidk: a toolkit to rapidly identify telomeric repeats from genomic datasets. Bioinformatics 41, btaf049 (2025).

121. A. Heger, C. Webber, M. Goodson, C. P. Ponting, G. Lunter, GAT: a simulation framework for testing the association of genomic intervals. Bioinformatics 29, 2046–2048 (2013).

122. A. R. Quinlan, I. M. Hall, BEDTools: a flexible suite of utilities for comparing genomic features. Bioinformatics 26, 841–842 (2010).

123. B. Gel, A. Díez-Villanueva, E. Serra, M. Buschbeck, M. A. Peinado, R. Malinverni, regioneR: an R/Bioconductor package for the association analysis of genomic regions based on permutation tests. Bioinformatics 32, 289–291 (2016).

124. M. Lawrence, W. Huber, H. Pagès, P. Aboyoun, M. Carlson, R. Gentleman, M. T. Morgan, V. J. Carey, Software for computing and annotating genomic ranges. PLOS Comput Biol 9, e1003118 (2013).

125. H. Išerić, C. Alkan, F. Hach, I. Numanagić, Fast characterization of segmental duplication structure in multiple genome assemblies. Algorithms Mol Biol 17, 4 (2022).

126. G. Benson, Tandem repeats finder: a program to analyze DNA sequences. Nucleic Acids Res 27, 573–580 (1999).

127. W. Shen, B. Sipos, L. Zhao, SeqKit2: A Swiss army knife for sequence and alignment processing. iMeta 3, e191 (2024).

128. P. Rice, I. Longden, A. Bleasby, EMBOSS: The European molecular biology open software suite. Trends Genet 16, 276–277 (2000).

129. M. Steinegger, J. Söding, MMseqs2 enables sensitive protein sequence searching for the analysis of massive data sets. Nat Biotechnol 35, 1026–1028 (2017).

130. T. Baril, D. Croll, MycoMobilome: a community-focused non-redundant database of transposable element consensus sequences for the fungal kingdom. NAR Genom Bioinform 8, lqag026 (2026).

131. H. Quesneville, C. M. Bergman, O. Andrieu, D. Autard, D. Nouaud, M. Ashburner, D. Anxolabehere, Combined evidence annotation of transposable elements in genome sequences. PLOS Comput Biol 1, e22 (2005).

132. T. Flutre, E. Duprat, C. Feuillet, H. Quesneville, Considering transposable element diversification in de novo annotation approaches. PLOS ONE 6, e16526 (2011).

133. C. Hoede, S. Arnoux, M. Moisset, T. Chaumier, O. Inizan, V. Jamilloux, H. Quesneville, PASTEC: An automatic transposable element classification tool. PLOS ONE 9, e91929 (2014).

134. K. Katoh, K. Misawa, K. Kuma, T. Miyata, MAFFT: a novel method for rapid multiple sequence alignment based on fast Fourier transform. Nucleic Acids Res 30, 3059–3066 (2002).

135. J. L. Steenwyk, T. J. B. Iii, Y. Li, X.-X. Shen, A. Rokas, ClipKIT: A multiple sequence alignment trimming software for accurate phylogenomic inference. PLoS Biol 18, e3001007 (2020).

136. B. Q. Minh, H. A. Schmidt, O. Chernomor, D. Schrempf, M. D. Woodhams, A. von Haeseler, R. Lanfear, IQ-TREE 2: New models and efficient methods for phylogenetic inference in the genomic era. Mol Biol Evol 37, 1530–1534 (2020).

137. G. Yu, D. K. Smith, H. Zhu, Y. Guan, T. T.-Y. Lam, ggtree: an r package for visualization and annotation of phylogenetic trees with their covariates and other associated data. Methods Ecol Evol 8, 28–36 (2017).

138. B. Gel, E. Serra, karyoploteR: an R/Bioconductor package to plot customizable genomes displaying arbitrary data. Bioinformatics 33, 3088–3090 (2017).

139. F. K. Mendes, D. Vanderpool, B. Fulton, M. W. Hahn, CAFE 5 models variation in evolutionary rates among gene families. Bioinformatics 36, 5516–5518 (2021).

140. A. Alexa, J. Rahnenfuhrer, topGO, version 2.56.0 (2025); http://bioconductor.org/packages/topGO/.

