## Supplementary Figures for "Transposable elements underlie chromosomal fusions and fissions in a highly species-rich group of butterflies"

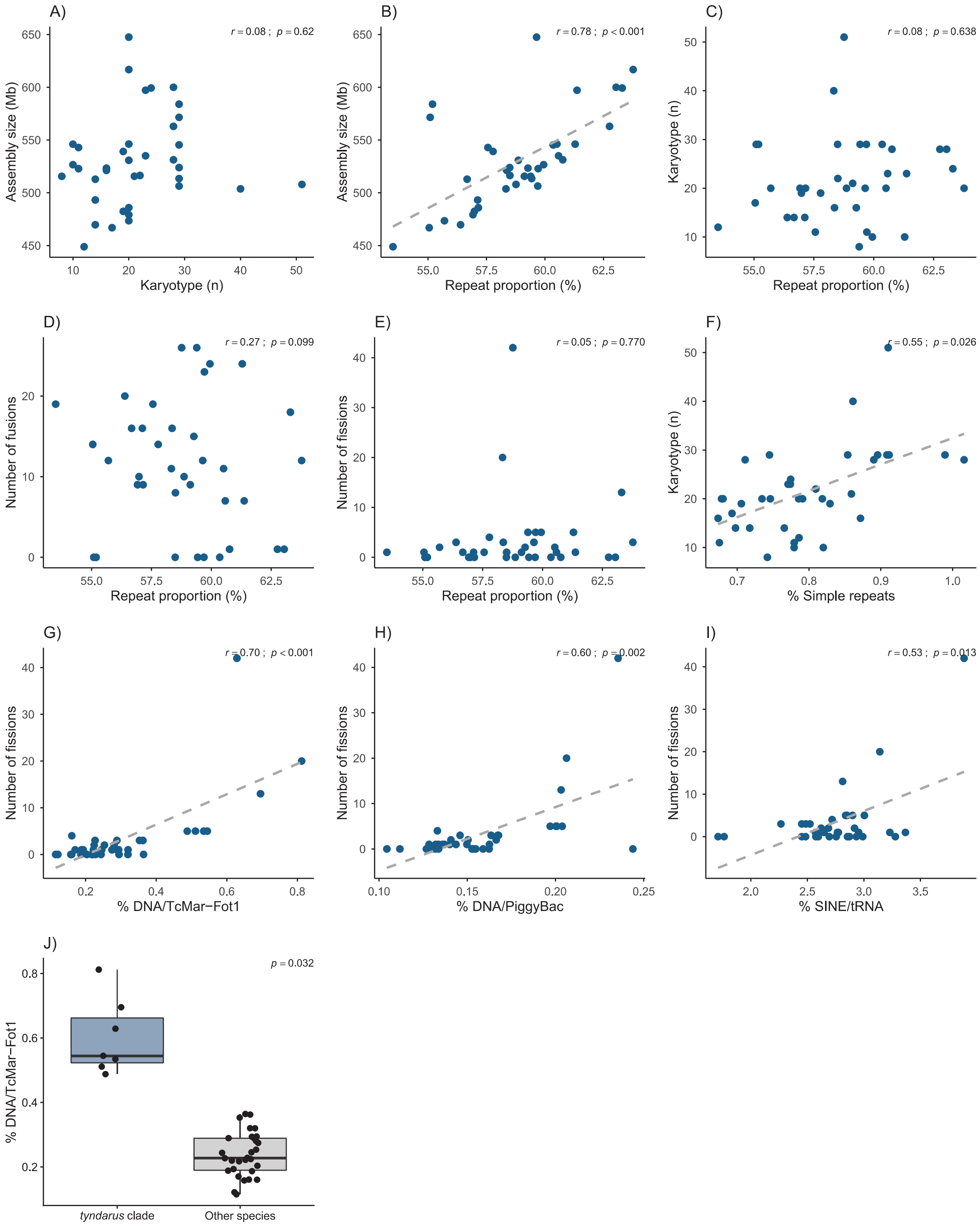


**Fig. S1.** Correlations between genomic features among the 37 *Erebia* reference genomes (including assembly size, karyotype, overall repeat proportion, number of fusions and fissions along the phylogenetic branches leading to extant species, and genomic proportions of simple repeats, DNA/TcMar-Fot1, DNA/PiggyBac and SINE/tRNA). The dashed grey lines represent significant Phylogenetic Generalized Least Square Regressions. The last plot represents a significant phylogenetic ANOVA between the *tyndarus* clade and other species in terms of genomic proportion of DNA/ TcMar-Fot1.

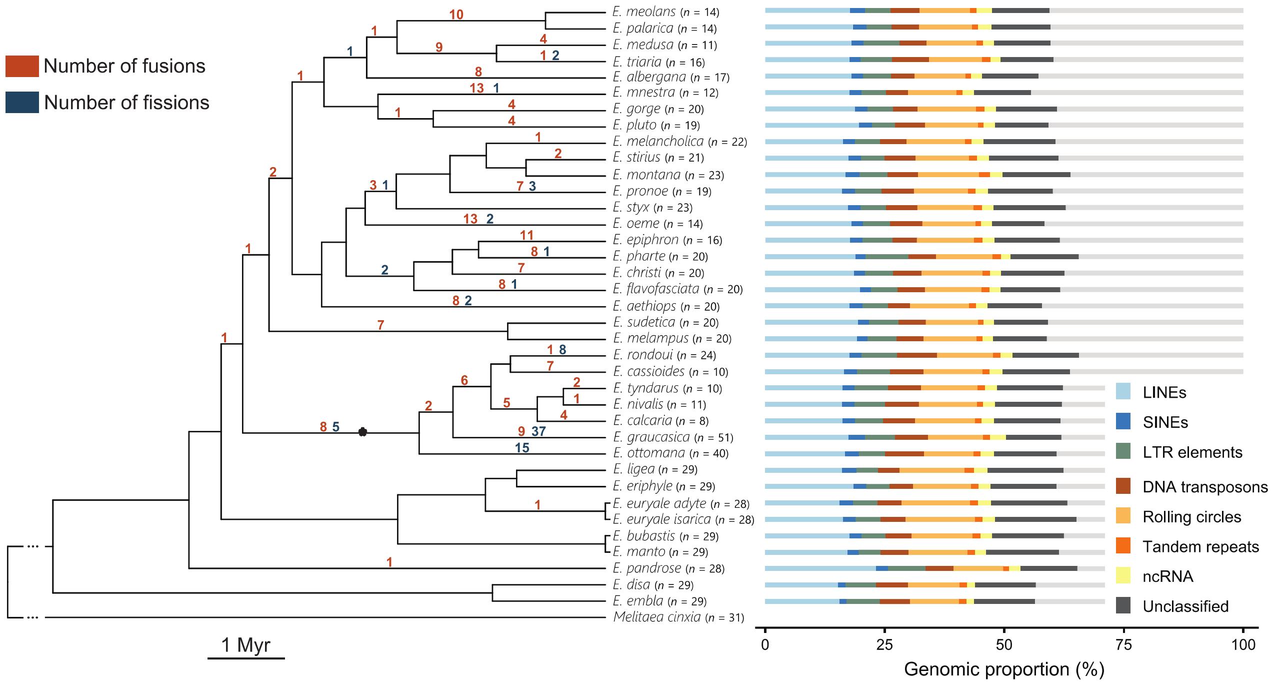


**Fig. S2.** Number of chromosomal fusions and fissions along the *Erebia* phylogeny and repeat proportions for each extant species. The star represents the branch of the *tyndarus* clade, i.e. the youngest, most species rich and karyotypically diverse *Erebia* clade. Haploid karyotype is indicated for each species, excluding the W chromosome. Repeat proportions in *Erebia* are calculated based on a genus-specific repeat library. LINEs = long interspersed nuclear elements, LTRs = long terminal repeats, SINEs = short interspersed nuclear elements, and ncRNAs = non-coding RNA.

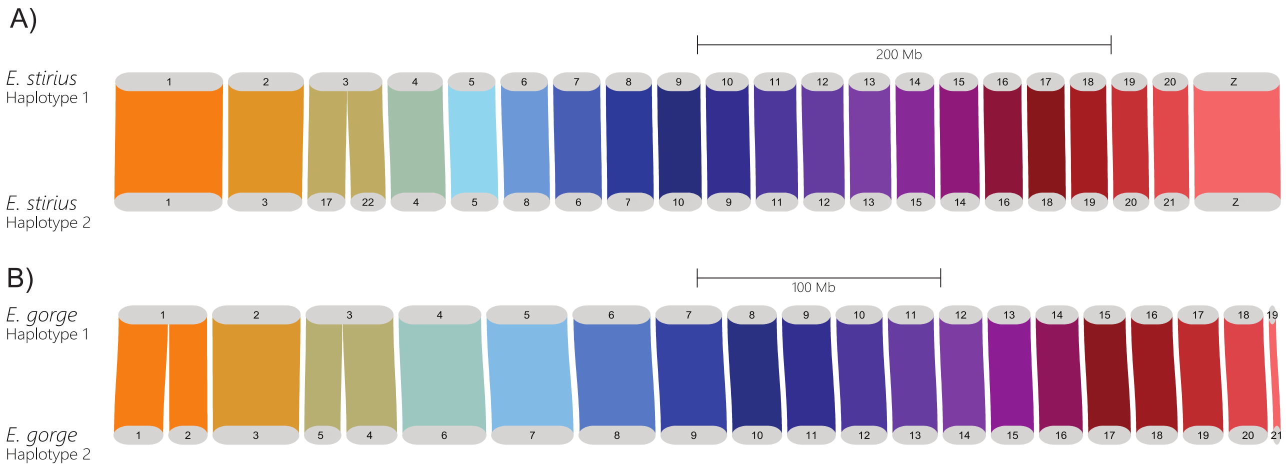


**Fig. S3.** Synteny between the two haplotypes of phased reference genomes generated from individuals heterozygous for chromosomal fusions. Chromosome 3 results from a fusion in *E. stirius* (A) and chromosomes 1 and 3 result from fusions in *E. gorge* (B). Note that the chromosomes are numbered according to size and do not represent homology between the two species. The reference genome of *E. gorge* was generated from a female, so sex chromosomes are not depicted.

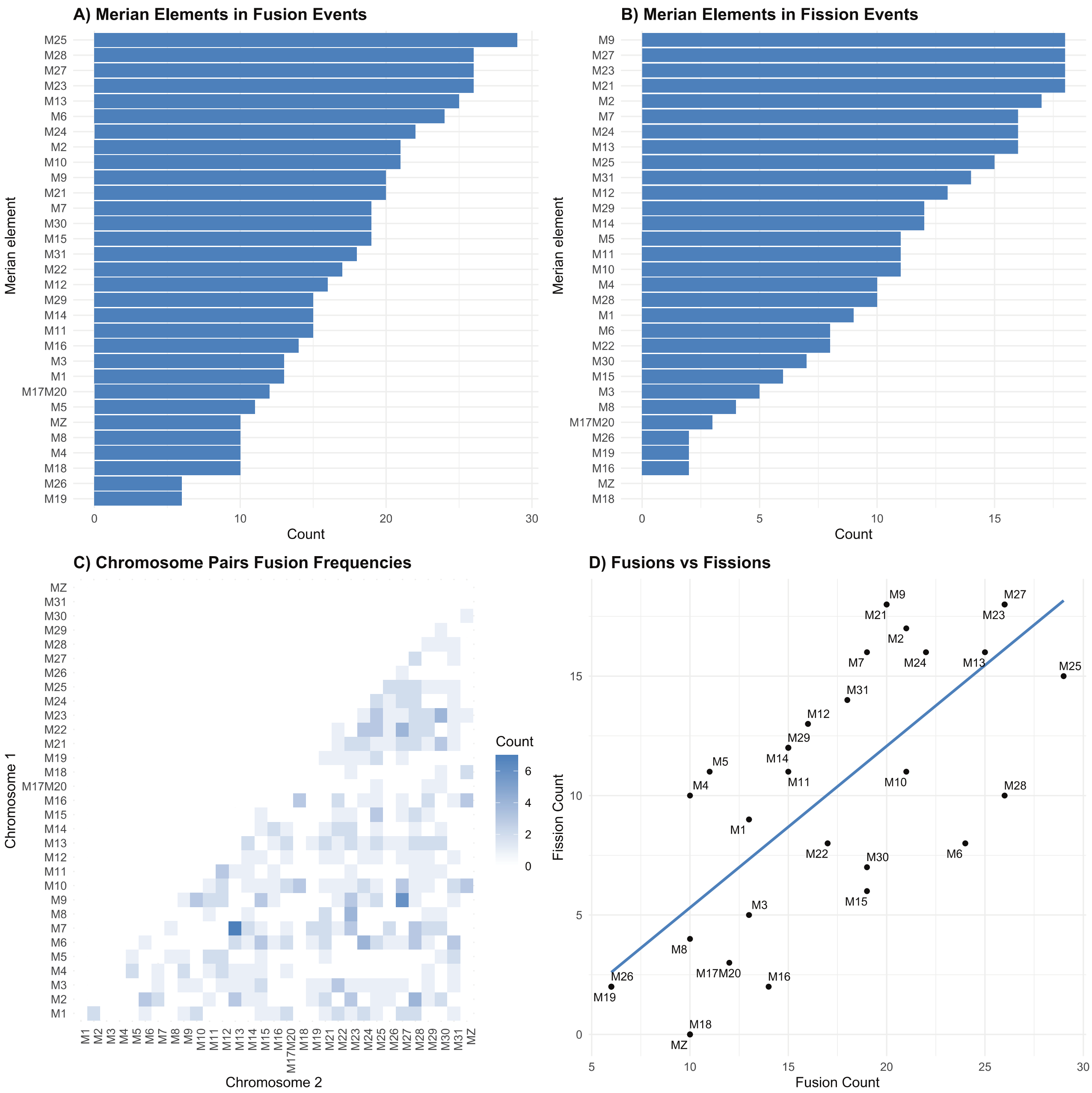


**Fig. S4.** Ancestral linkage groups in Lepidoptera (i.e. Merian elements, M) and their tendency to be involved in chromosomal fusions and fissions in *Erebia* butterflies. (A) Number of chromosomal fusions detected for each Merian element. (B) Number of chromosomal fissions detected for each Merian element. (C) Number of fusions detected between each Merian element pair. (D) Positive correlation between the number of fusions and the number of fissions detected for each Merian element. We grouped M17 and M20 together as they represent a very ancient fusion (Wright et al. 2024).

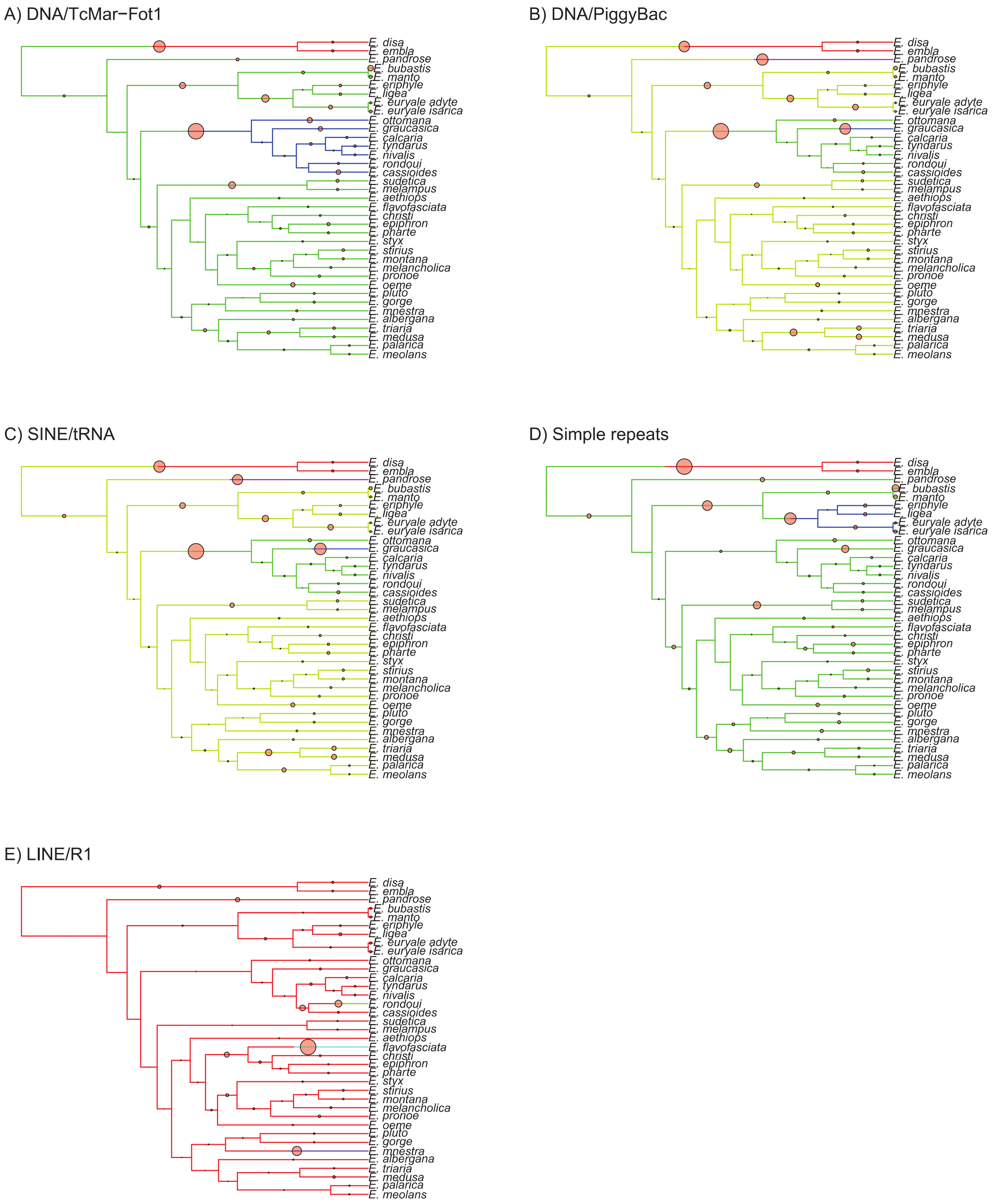


**Fig. S5.** Results of bayou analyses (Bayesian reversible-jump multi-optima OU models) for DNA/TcMar-Fot1, DNA/PiggyBac, SINE/tRNA, Simple repeats and LINE/R1. Colour changes along the branches represent significant optima shifts (Posterior Probability (PP) > 0.3, red circles proportional to PP).

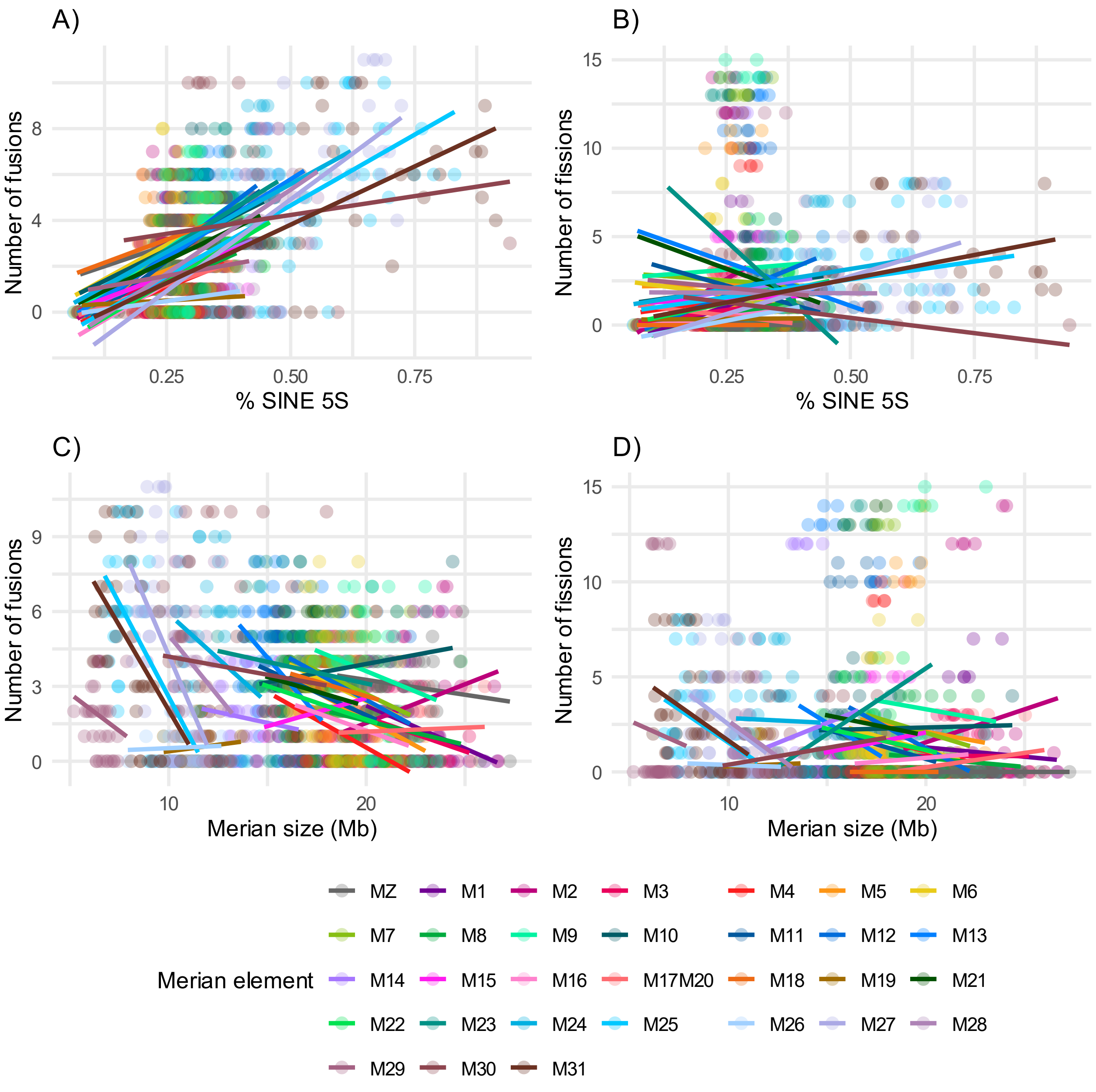


**Fig. S6.** Correlations, separated per Merian elements, between number of fusions and fissions along the phylogenetic branches leading to extant species and genomic proportions of SINE/5S and chromosome size. M17 and M20 are depicted and analysed together as they represent a very ancient fusion (Wright et al. 2024).

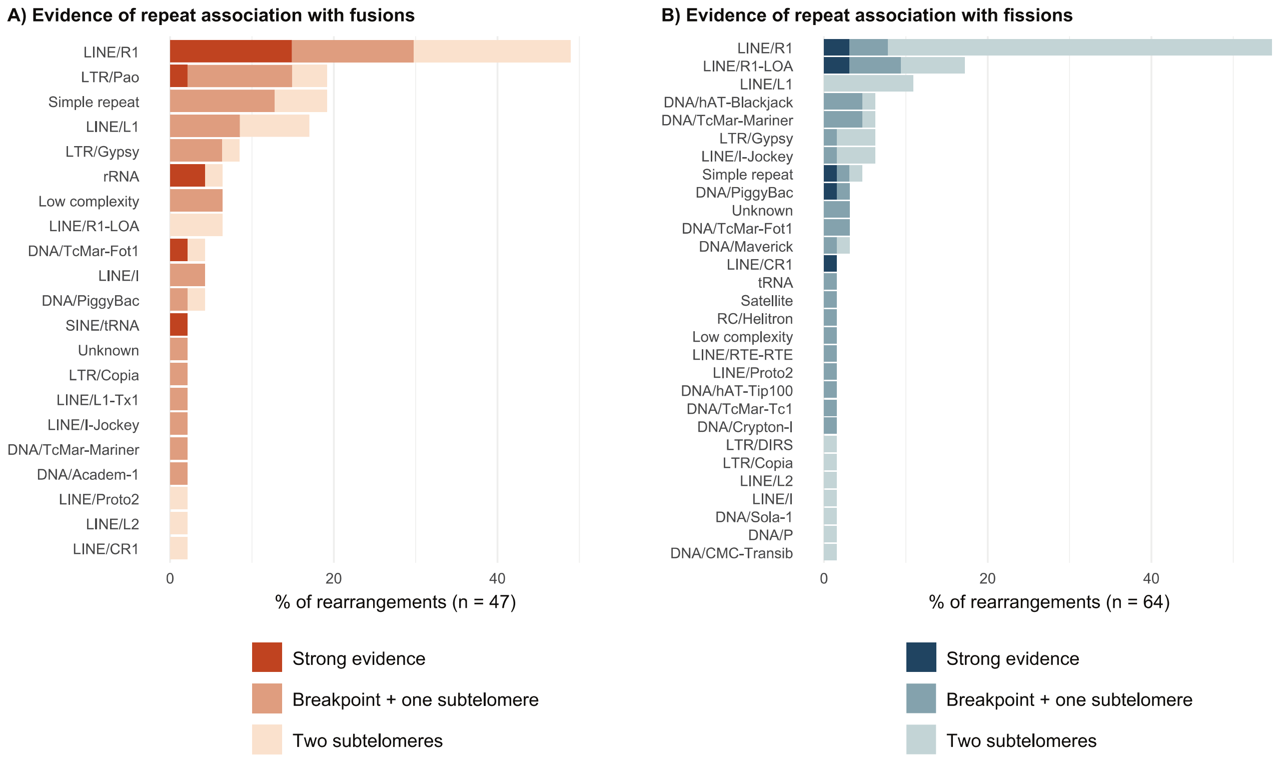


**Fig. S7.** Number of fusions (A) and fissions (B) with which each repeat family is associated, based on the enrichment calculated with the software *regioneR*. Evidence of repeat association is divided into strong evidence, i.e. repeats enriched at the breakpoint and both corresponding homologous regions in a closely related species with the unfused state, or supporting evidence when repeats are enriched either at the breakpoint and one homologous region, or at both homologous regions.

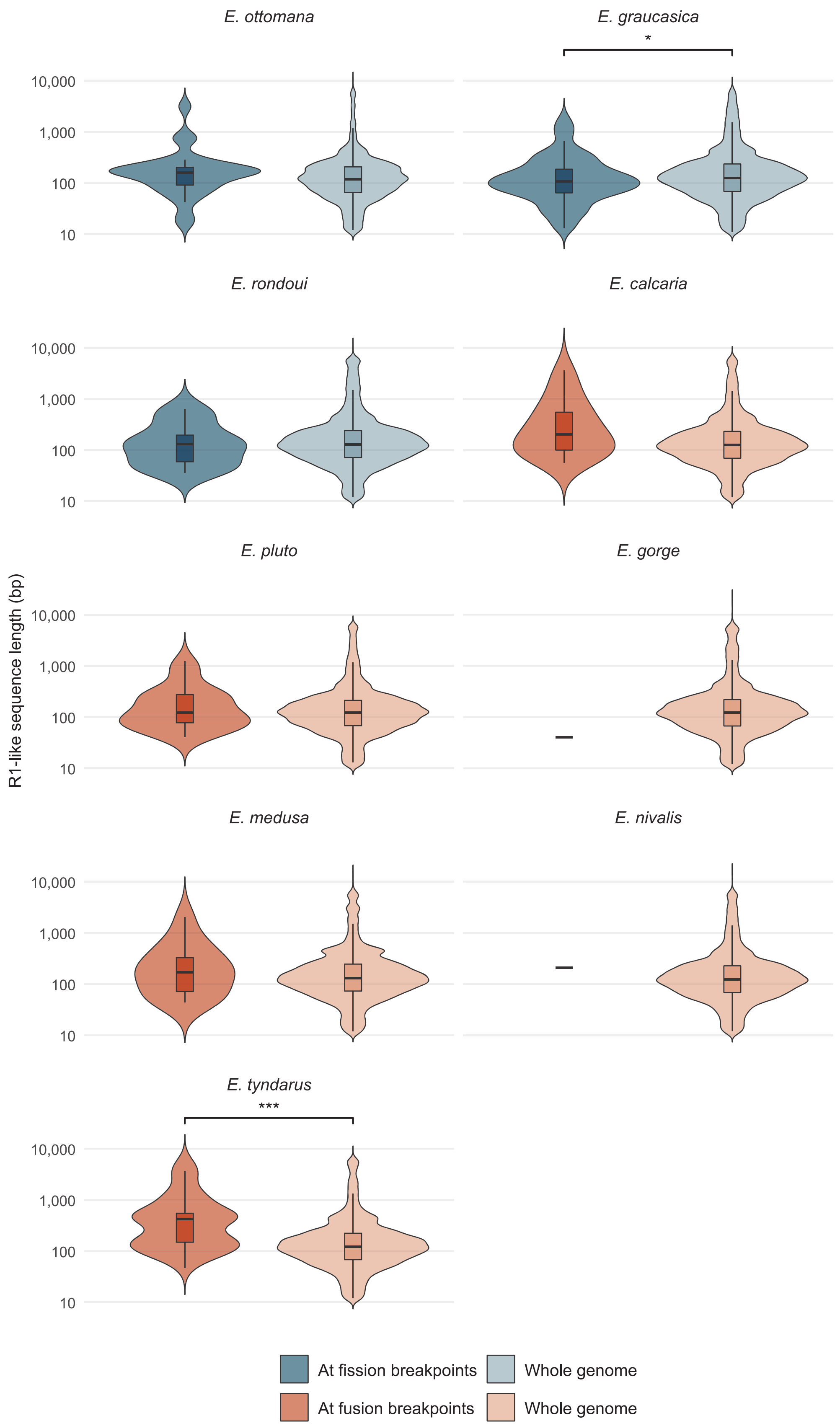


**Fig. S8.** Sequence length of R1-like elements at fusion and fission breakpoints compared to genome-wide for the species without strong evidence for R1-like elements association with rearrangements (permutation test: *** $p < 0.001$, ** $p < 0.01$, * $p < 0.05$).

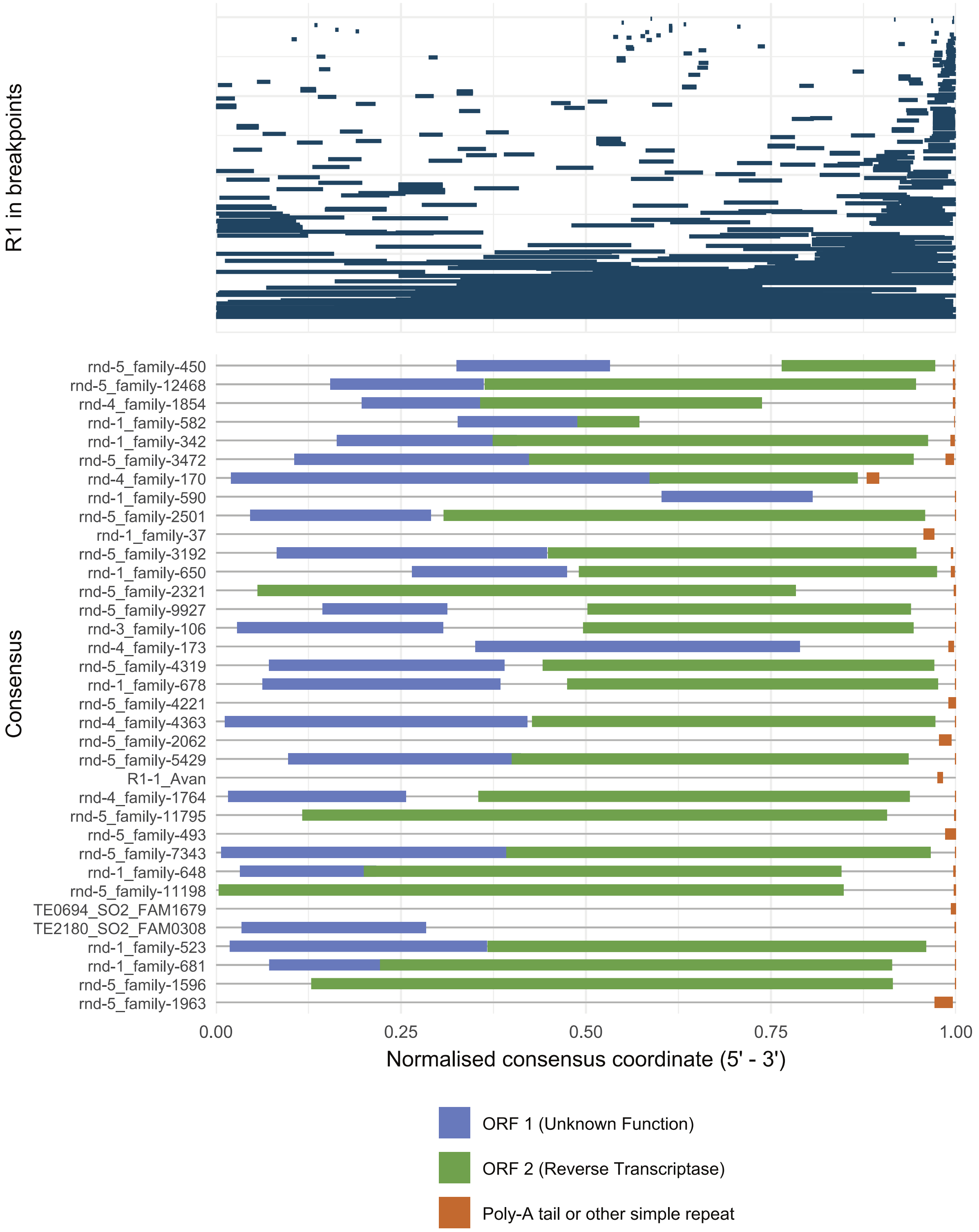


**Fig. S9.** Top: R1-like fragments annotated at fusion and fission breakpoints, for consensus sequences that could be oriented (5’ – 3’) given the presence of a poly-A tail, or other short tandem repeat, at the 3’ end. The x-axis indicates normalised start and end coordinates for each element relative to its consensus. Fragments are organised from shortest to longest. Bottom: For the same R1-like element consensus sequences, normalised coordinates of the two canonical open-reading frames (ORF) and the poly-A tail. Consensus sequences are organized from most abundant at breakpoints to least abundant.

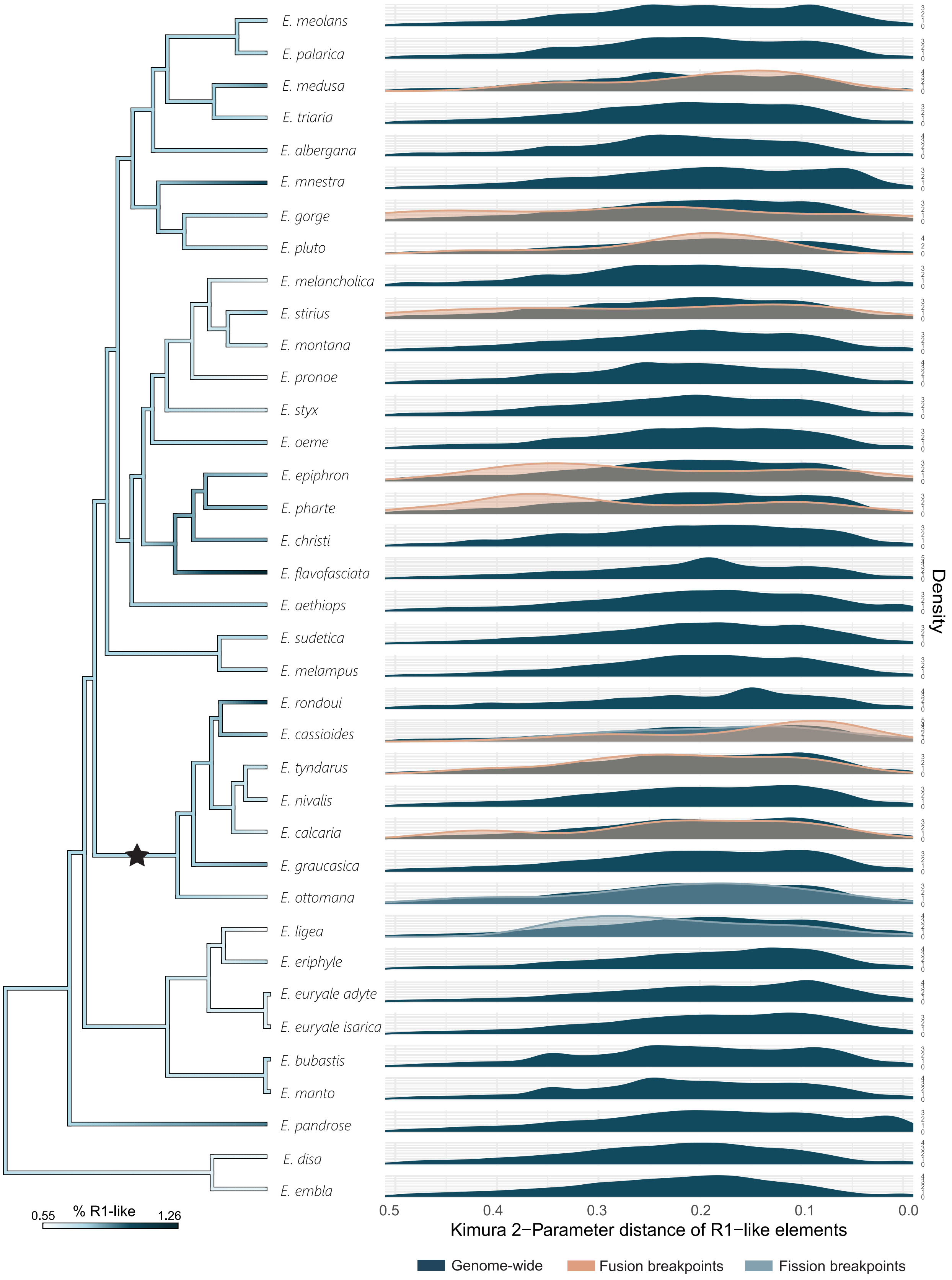


**Fig. S10.** R1-like genomic proportion across the *Erebia* phylogeny displayed as colour gradient (left) and density plots of Kimura 2-parameter distance distribution of LINE R1-like in each species and at fusion and fission breakpoints (right). Note that the Kimura axis represents an inverted scale with more similar repeats on the right.

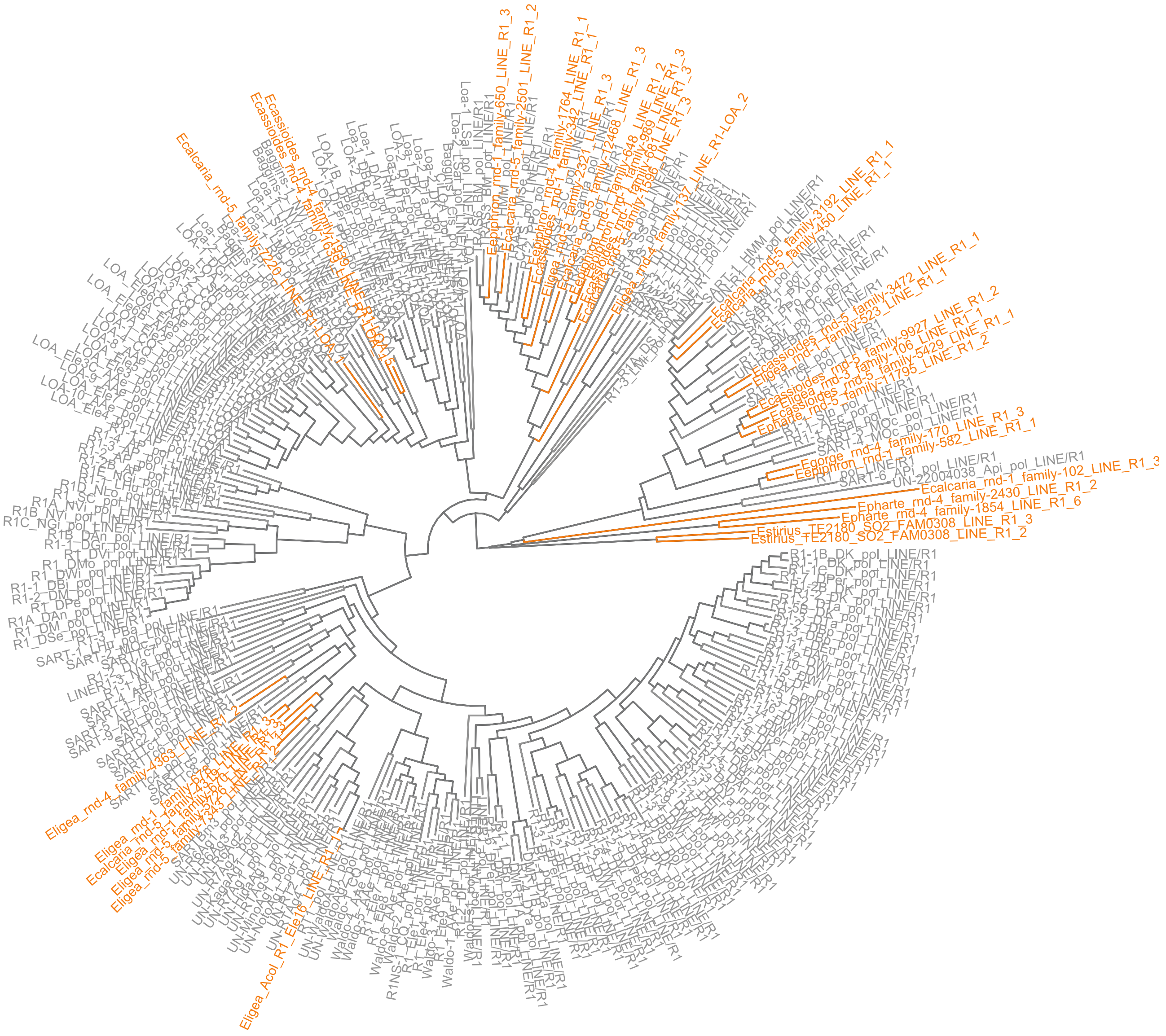


**Fig. S11.** Phylogenetic tree of LINE R1-like elements across *Erebia*. The elements found at fusion or fission breakpoints are shown in orange.

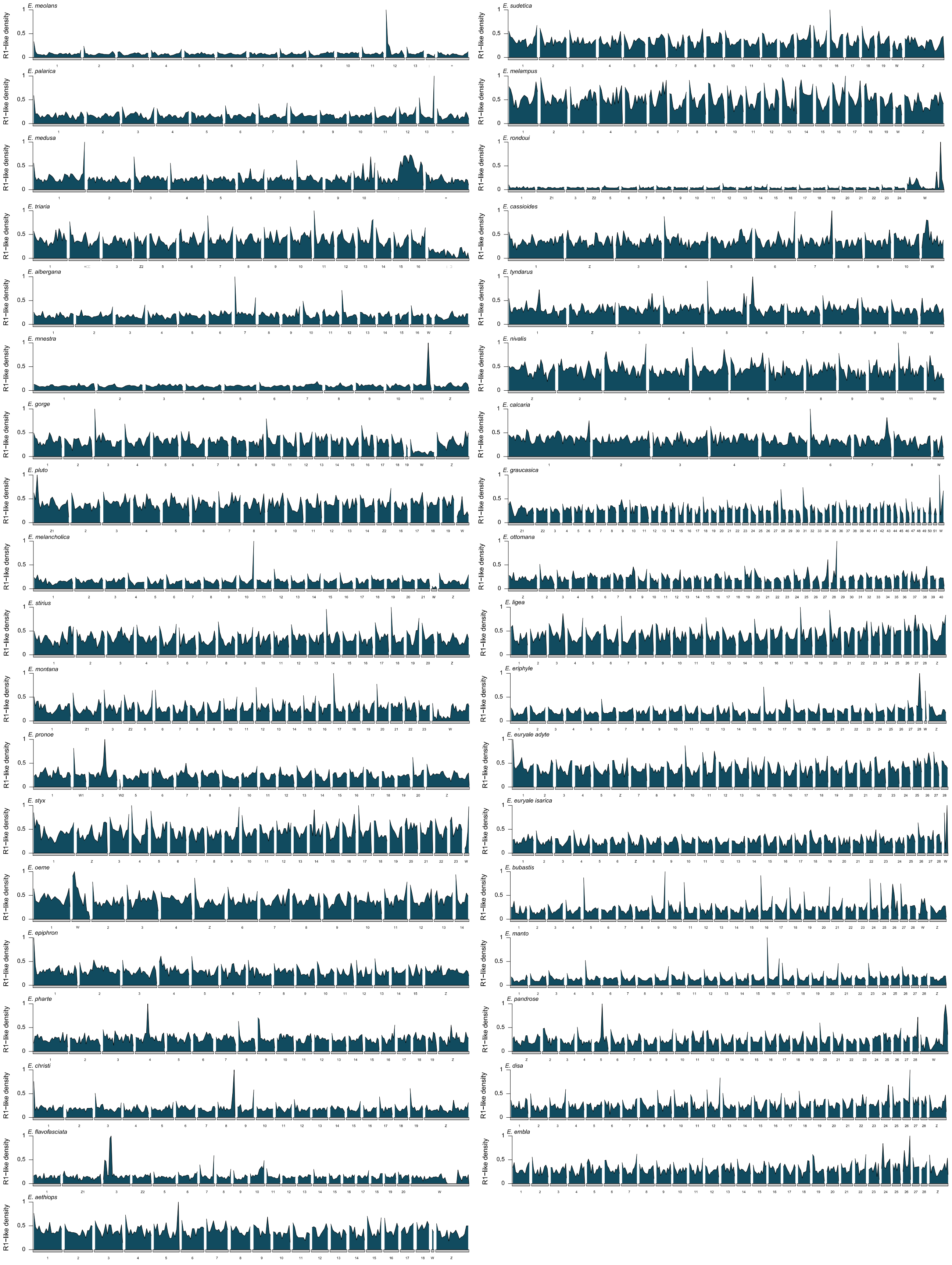


**Fig. S12.** LINE R1-like genomic density distribution across the 37 *Erebia* reference genomes. Y-axis is the normalized (from zero to one in each species) number of R1-like elements in a 200 kb genomics window.

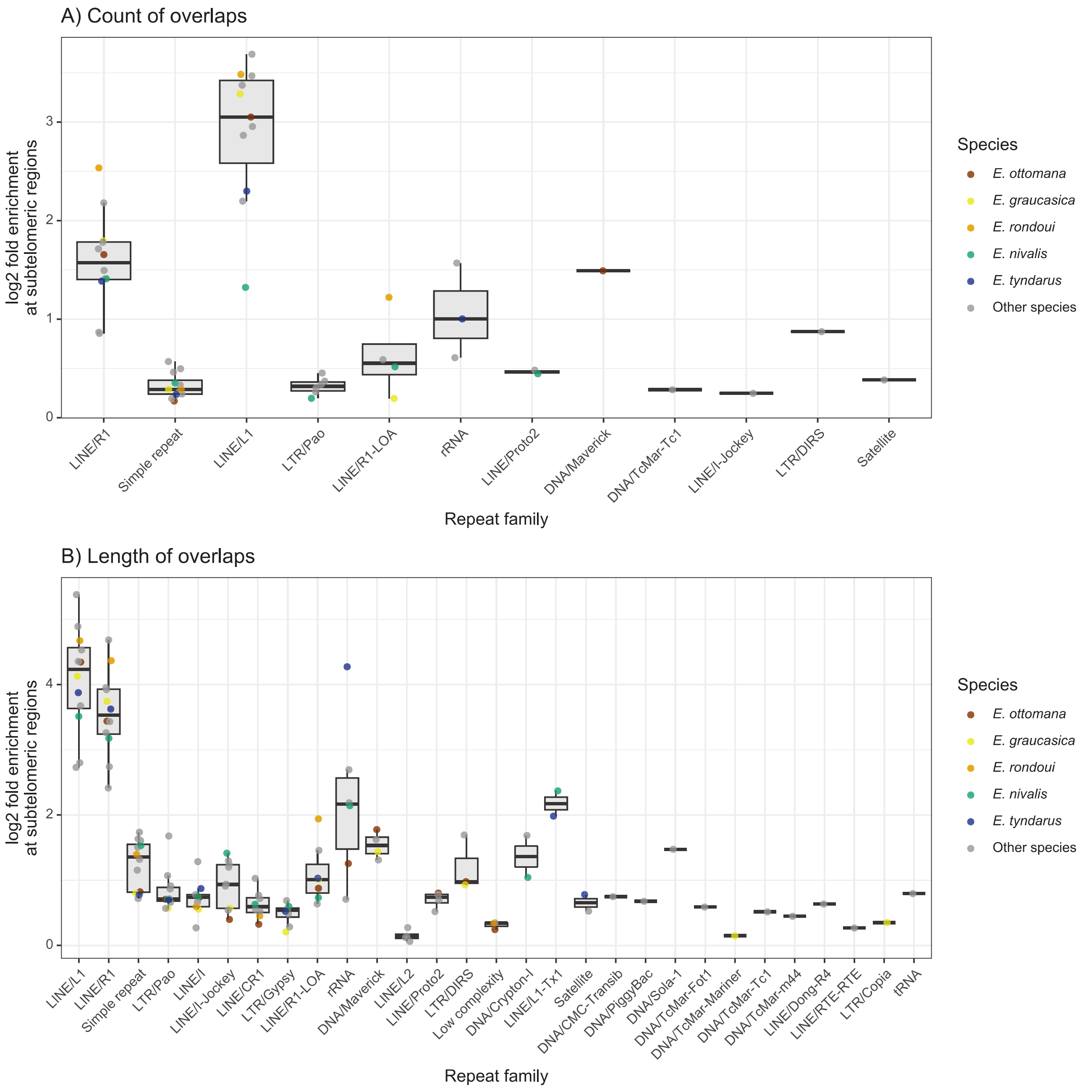


**Fig. S13.** Enrichment of repeat families at subtelomeric regions in species for which repeat enrichment at rearrangement-associated subtelomeres was tested. Bars show the log2 fold enrichment relative to genome-wide levels for repeat families with significant subtelomeric enrichment ($p < 0.05$), based on the number of overlaps (A) or the total length of overlaps (B). Only species and repeat families with significant enrichment are shown. Species not belonging to the *tyndarus* clade are presented in grey, species belonging to the *tyndarus* clade in different colours.

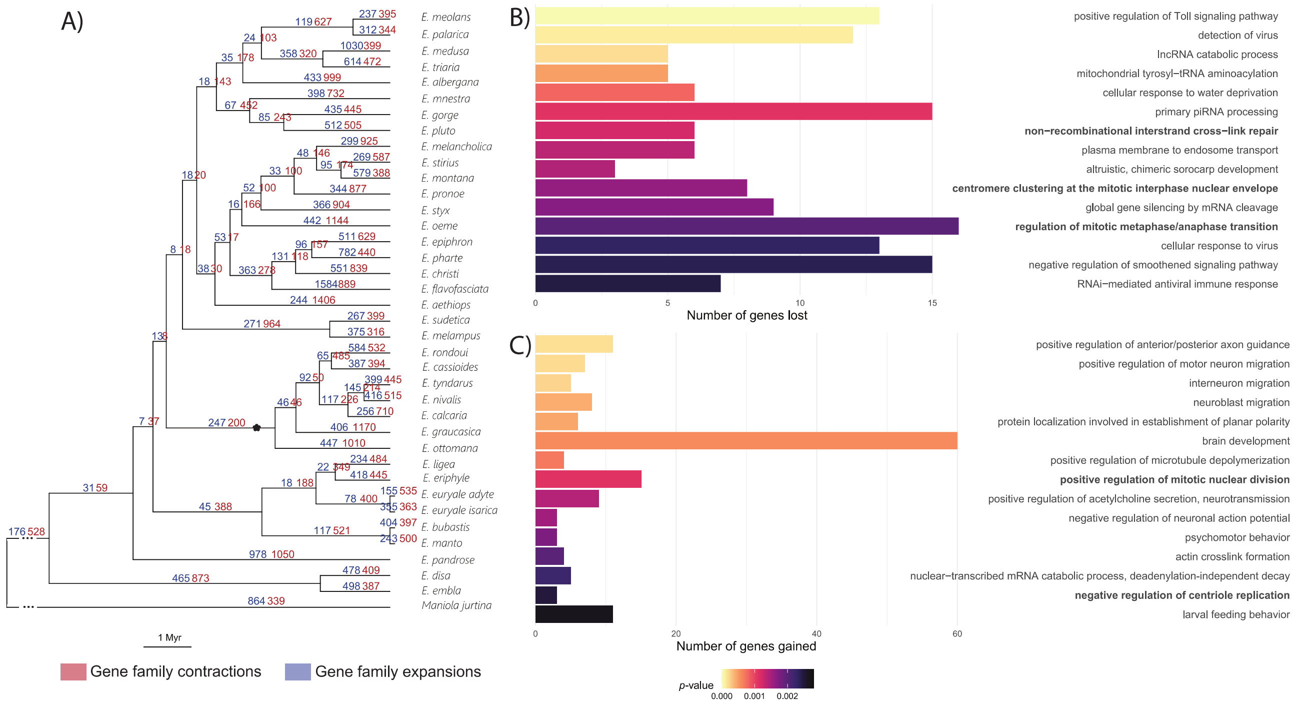


**Fig. S14.** Gene family expansions and contractions along the *Erebia* phylogeny. The node separating the karyotypically variable *tyndarus* clade is depicted with a star (A). GO term enrichment analysis on the gene family contractions (B) and expansions (C) at the base of the *tyndarus* clade. We only report GO terms that are not enriched in the gene family expansions and contractions at the base of the karyotypically conserved *ligea* clade. In bold are GO terms potentially involved in chromosomal rearrangements. The full list of enriched GO terms can be found in Tables S17 and S18.

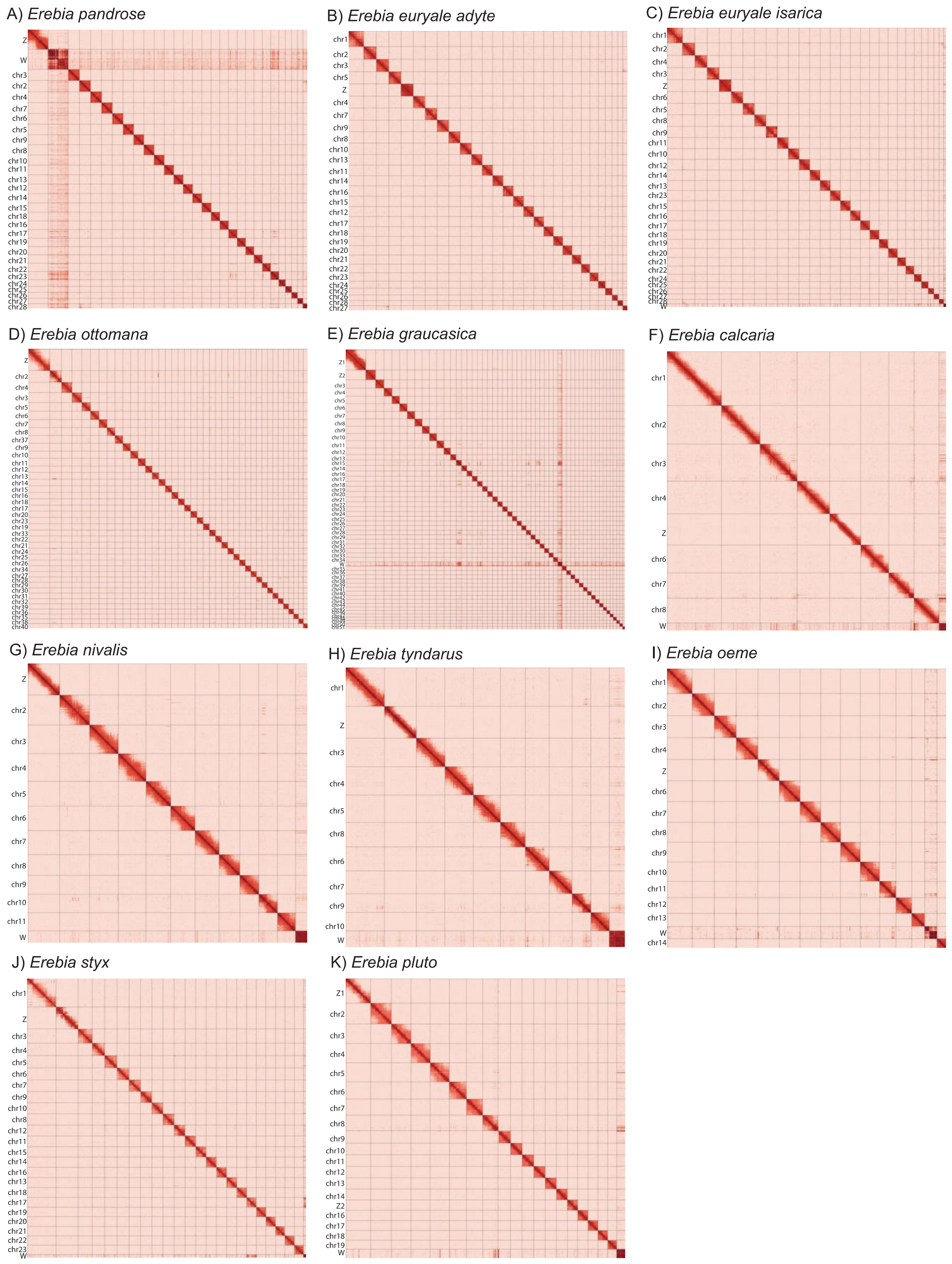


**Fig. S15.** Final Hi-C contact maps of the 11 *Erebia* genome assemblies presented in the present manuscript. Assembled chromosomes are shown in order of size. The plot was generated using PretextSnapshot v.0.0.5 (https://github.com/sanger-tol/PretextSnapshot, with options -r 4000 -c 30).

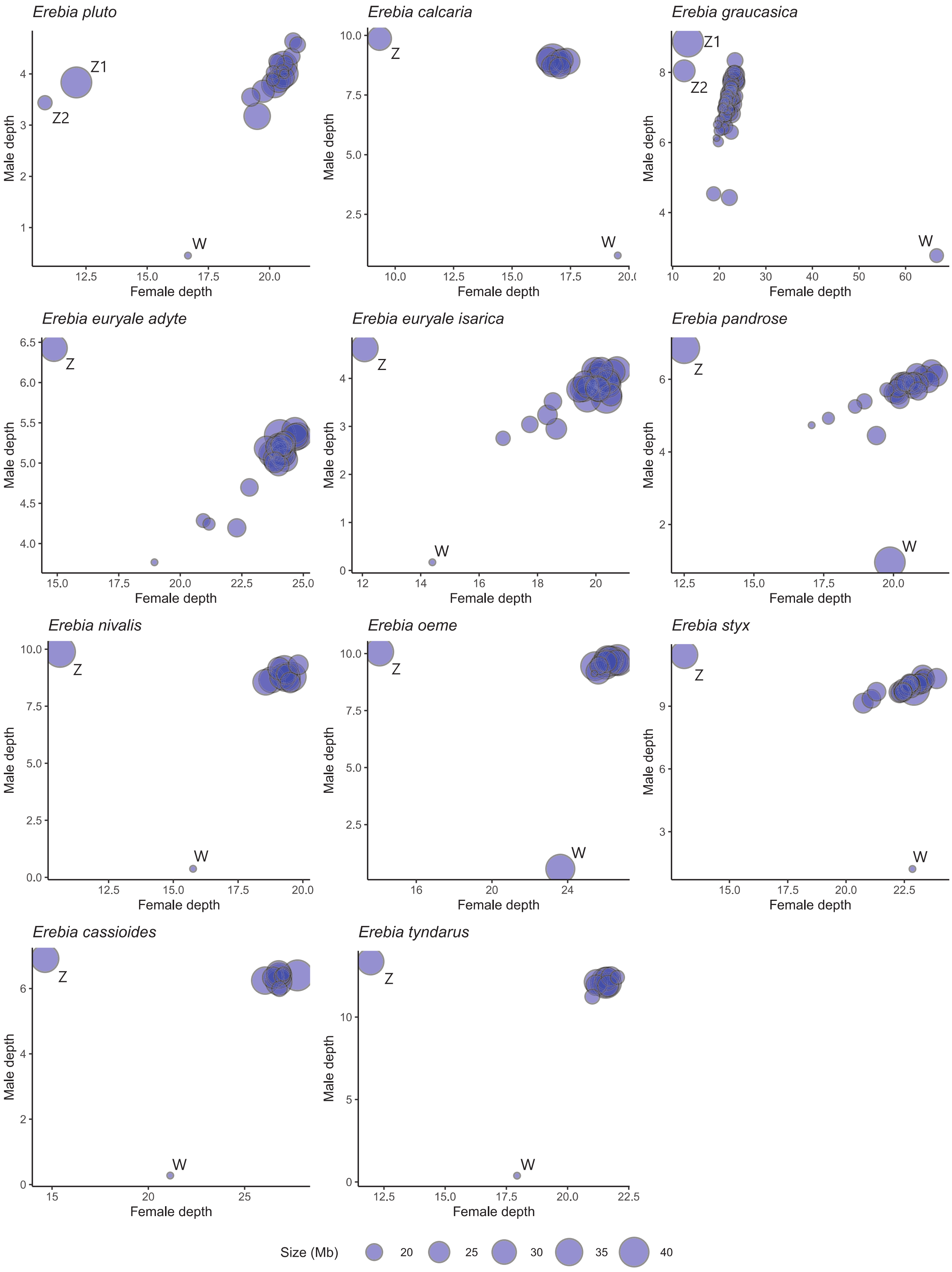


**Fig. S16.** Sequencing depth of HiFi reads (for female individuals) and Illumina reads (for male individuals) mapped back to the reference genome to identify Z and W sex chromosomes. Dot size is proportional to chromosome size.
